# Leaf-age specific flood resilience in *Arabidopsis thaliana* is determined by distinct processes contributing to post-submergence recovery

**DOI:** 10.1101/2025.10.16.682766

**Authors:** Tom Rankenberg, Maria Angelica Sanclemente, Hongtao Zhang, HACF Leeggangers, Muthanna Biddanda Devaiah, Janet Rong Chao, Hans van Veen, Frederica L. Theodoulou, Rashmi Sasidharan

## Abstract

While the molecular mechanisms mediating submergence tolerance have been extensively studied, those underpinning age-dependent resilience remain poorly characterized. In *Arabidopsis thaliana*, submergence elicits a leaf-age dependent phenotype in which senescence and death progress across an age gradient starting with older leaves. Here we sought to investigate the mechanisms mediating this observed differential flood resilience by interrogating leaf age-specific transcriptome and proteome changes during submergence and recovery. Following submergence, most age-dependent differences were in the magnitude or speed of transcript abundance changes, whereas qualitative leaf-age dependent responses were most apparent during recovery. This included a strong desiccation response in old leaves despite a stronger ABA-signaling response. Physiological measurements suggested that faster dehydration was facilitated by a combination of submergence-mediated reduction of ABA sensitivity and higher conductance in old leaves. We also observed a stronger induction in young shoot tissue of genes associated with endoplasmic reticulum (ER) stress and the unfolded protein response (UPR). Mutants disabled in the two UPR signaling branches were affected in new leaf formation and the ability to restore the proteome, but not in senescence, suggesting that young tissues activate ER stress recovery to permit continuation of growth. Of the mitochondrial membrane proteins differentially regulated in young leaves, loss of mitochondrial voltage-dependent anion channel function impacted submergence-recovery tolerance. Our data reveal multiple mechanisms underlying leaf age-dependent differential submergence recovery and demonstrate how tolerance is determined by an interplay between age related developmental traits and stress signaling pathways.

## Introduction

Complete submergence exposes the entire plant to the stress (Sasidharan et al., 2017). However, the distinct identity of plant tissues can cause substantial differences in stress sensing and responses. This variation is not only a consequence of differences in physical properties but can also be due to differences in the molecular regulation of stress responses, ultimately leading to differential stress resilience (Rankenberg et al., 2021). Previous research has demonstrated both shared and unique responses to hypoxia and submergence stress between different cell types (Mustroph et al., 2009). Waterlogging induces the expression of cell wall modifying enzymes, specifically in the root cortex leading to aerenchyma formation (Rajhi et al., 2011). Submergence can also elicit organ-specific responses (Van Veen et al., 2016). These studies compared cell types or organs that fulfill very different roles within a plant. However, variation in submergence responses can also be determined by plant age. In Arabidopsis, submergence tolerance declines during the transition from two-week old seedlings to five-week adult plants. This decrease in tolerance stemmed from a reduction in the ability of Ethylene Response Factor (ERFVIIs) and the NAC transcription factor ANAC017 to activate their downstream targets (Giuntoli et al., 2017; Bui et al., 2020). During submergence, the activity of the ERFVII RAP2.12 is limited by the trihelix transcription factor HYPOXIA RESPONSE ATTENUATOR1 (HRA1) (Giuntoli et al., 2014). HRA1 binds to RAP2.12 and reduces its activity as a transcriptional activator, which attenuates the induction of anaerobic metabolism by RAP2.12. The repression of RAP2.12 activity by HRA1 is strongest in young leaves and meristems, where it likely serves to limit the utilization of carbon resources during hypoxia. Limiting carbon consumption during hypoxia is not only beneficial to preserve energy for post-stress recovery but also prevents premature activation of metabolic adjustments to hypoxia, since meristems are already hypoxic under non-flooded conditions (Weits et al., 2014).

These studies have thus clearly demonstrated how differential regulation of principal hypoxia response regulators such as ERFVIIs, HRA1 and ANAC017 lead to age-associated changes in submergence tolerance. The decrease in flooding tolerance with age is observed for both whole plants and individual leaves. Changes in stress tolerance of tissues with age can simply be a consequence of changes in physiology but can also be an actively regulated process. Several specific transcription factors have recently been linked to changes in age-dependent stress tolerance (D’alessandro et al., 2018; Zhao et al., 2022) suggesting that age-dependent stress responses are often actively regulated at a molecular level.

A frequently observed pattern in age-dependent stress responses is a decrease in stress tolerance with leaf age (Rankenberg et al., 2021; Rankenberg et al., 2024). This can be caused by the inability of old leaves to properly respond to a stress (D’alessandro et al., 2018), or by the faster induction of stress-induced senescence in old leaves (Schippers, 2015). Actively allowing an old leaf to die by accelerating the onset of senescence while a young leaf remains alive can be beneficial to a stressed plant, as nutrients can be transported from dying old tissues to young tissues and meristems to fuel their survival (Yu et al., 2015). Prioritizing the survival of more expendable tissues to keep meristems alive could permit plant reproductive success during stressful conditions. Despite the observed age dependence of stress tolerance in many species, the underlying molecular mechanisms are underexplored. Most studies tend to neglect the age dimension, focusing instead on bulk tissues where such differences might be lost. Here we capitalized on the very clear age-directed senescence gradient induced in Arabidopsis rosettes upon complete submergence. Furthermore, sequentially experiencing flooding followed by re-aeration stress results in a set of confounding effects determining the success of individual plant organs. How organ age and confounding effects intersect to determine performance, and what the limiting/causative factors are remains unclear.

To identify the processes and regulators that play a role in this age-dependent response to submergence stress, we characterized and compared the transcriptomes of an old and a young leaf of Arabidopsis during and after complete submergence. Both the submergence and post-submergence phases triggered significant changes in old and young leaf transcriptomes. Shared transcriptome responses to submergence between old and young leaves included the general repression of translation and growth. In old leaves, genes associated with leaf senescence exhibited increased expression during both the submergence and post-submergence phase. However, prolonged submergence treatment eventually also induced the expression of senescence-associated genes in young leaves. Although a strong correlation was observed between old and young leaf responses during the submergence phase, this was largely lost during recovery. The leaf -age dependent divergence of transcriptomic responses during recovery was consistent with the visual onset of stress between leaves during this phase. To further validate findings from the transcriptomics data we thus focussed on leaf-age dependent processes evident during <u>recovery</u>: a strong desiccation transcriptome signature in old leaves and a stronger induction of endoplasmic reticulum (ER) stress genes in young leaves. Surprisingly, faster dehydration in old leaves could not be attributed to changes in ABA biosynthesis, stomatal density, aperture or cuticle thickness. Instead our results suggest that a combination of decreased ABA sensitivity and higher leaf conductance make older leaves dehydrate faster following desubmergence. Consistent with our transcriptomics findings, ER stress sensing and signaling is important for submergence recovery. Mutants for components of ER stress sensing were compromised in submergence survival. Importantly, mutant phenotypes deviated in terms of ‘young leaf’ attributes i.e. new leaf formation and not submergence-induced senescence. ER stress responsive genes were also preferentially induced in young leaves during recovery. This was confirmed by comparison of the transcriptome and proteomes of mutants disabled in ER stress sensing and signaling. This global assessment also revealed the preferential upregulation in young leaves of proteins contributing to maintainence of cellular homeostasis and faster recovery. This included a group of mitochondrial proteins, the voltage dependent anion channels, that have recently been implicated in mediating mitophagy (Ma et al., 2025). Overall our data provides a comprehensive insight into molecular pathways mediating leaf-age dependent differential submergence sensing and recovery.

## Results

### Submerged Arabidopsis rosettes exhibit age-dependent sequential leaf death

Complete submergence in darkness (hereafter submergence) of Arabidopsis Col-0 plants for varying durations triggered a sequential leaf death pattern (Fig. 1A). Visible senescence and leaf death was initiated in the oldest leaves and progressed towards younger leaves, consistent with our previous observations (Rankenberg et al., 2024). Plants recovering from sub-lethal submergence treatments showed further deterioration following desubmergence, with senescence and leaf dehydration in the post-submergence phase being more severe in old and intermediate leaves than in young leaves. The age-dependent effects of the submergence and post-submergence phases led to age-dependent leaf death: the old leaves died faster than the young leaves (Fig. 1B). To characterize the molecular responses and potential regulatory genes and processes underlying this differential age effect, a mRNA-seq approach was used. Blades of leaf three and seven of a 10-leaf Arabidopsis plant (Fig. 1A) were sampled at eight timepoints (Fig. 1C). Leaf three was chosen as a representative ‘old’ leaf. It resembles most other rosette leaves, apart from leaves one and two, which have a rounder shape and die faster than the rest (Fig. 1C). Leaf number seven was the representative ‘young’ leaf. It is the youngest leaf at this growth stage that has an easily distinguishable petiole. Despite their physiological similarities, these two leaves died at different rates following submergence (Fig. 1B). Sampling for mRNA-seq occurred before submergence (t0_00), after two (t2_00), four (t4_00), and six (t6_00) days of submergence. Leaf samples were also harvested after four days of submergence at one (t4_01), three (t4_03), six (t4_06), and twenty-four (t4_24) hours in the light after desubmergence (Fig. 1C). Only the young leaf was sampled at the six days submergence timepoint, as the old leaf was often dead at this time. Differential gene expression was calculated at: (i) each submergence timepoint compared to the non-submerged plants (the submergence effect); (ii) each post-submergence phase time point relative to four days submerged plants (the recovery effect); (iii) each post-submergence timepoint compared to the non-submerged plants (the full effect). The first two approaches allowed us to investigate the specific response to the submergence and post-submergence phases without including the effect of the sequential exposure to these phases.

**Figure 1.**
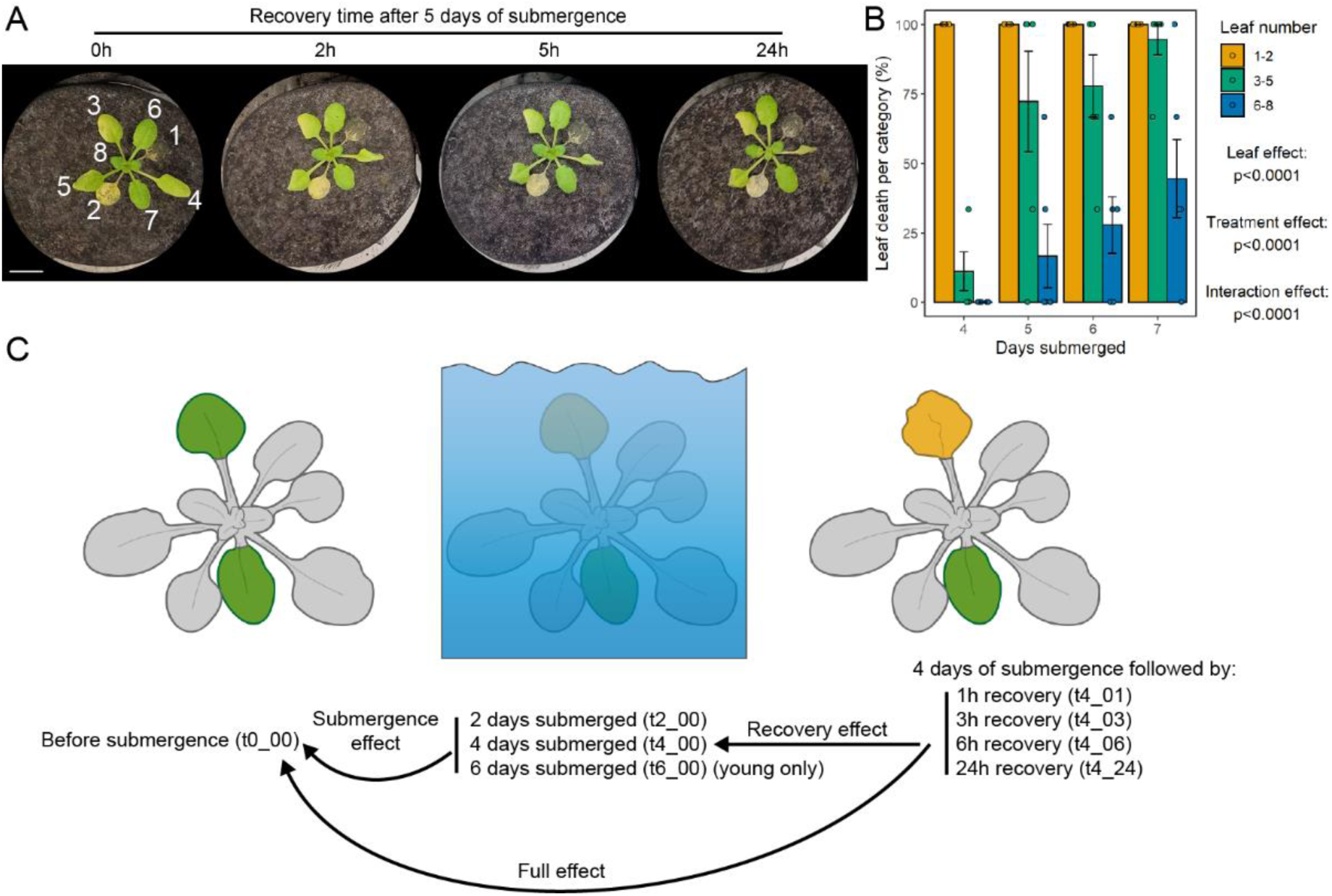
| Resilience to the submergence and post-submergence stress decreases with leaf age A) Representative images of wild-type Arabidopsis plants (accession Col-0) recovering from five days of dark submergence and showing age-dependent leaf damage and death. Images were taken at the recovery time points indicated. Numbers in the first image (0h recovery) indicate leaf numbers. Scale bar indicates 1 cm. B) Submergence causes age-dependent leaf death. Leaf death per age category was scored three days after desubmergence after the indicated submergence durations. Significance was determined by two-way ANOVA (leaf age*time). Leaf numbers are as indicated in Fig. 1A. C) Overview of the mRNA-seq experimental design and treatment comparisons. Laminas of old (leaf three) and young (leaf seven) leaves of 10-leaf-stage Arabidopsis plants were harvested before submergence and after the indicated treatments. Gene expression was calculated for each submergence timepoint relative to non-submerged conditions (submergence effect), for the post-submergence timepoints relative to the last submerged timepoint before desubmergence (recovery effect), and for the post-submergence timepoints relative to the non-submerged conditions (full effect).

### The submergence and post-submergence phases together have a stronger effect on the transcriptome of old leaves

Multidimensional scaling of the 2000 most variable genes showed that transcriptomes of old and young leaves were relatively similar before submergence (t0_00, Fig. 2A). However, the submergence time points (t2_00, t4_00, and t6_00) all clustered far away from the pre-submergence time points, suggesting massive transcriptional reprogramming during submergence in both old (circles) and young leaves (triangles). The post-submergence time points (t4_01, t4_03, t4_06, and t4_24) of old leaves revealed a second phase of transcriptional reprogramming. At 24h after desubmergence, old leaf transcriptomes were still far removed from the t0_00 time point. This was in contrast with the pattern of young leaf post-submergence timepoints, where the last post-submergence timepoint clustered very close to the t0_00 timepoint, suggesting more rapid recovery. The number of differentially expressed genes (DEGs) detected for old and young leaves was consistent with the MDS plots. DEG numbers were similar at submergence time points but diverged during recovery with lower numbers in young than in old leaves (Supplemental Fig. S1A; Supplemental Table S1). These patterns were consistent with observed leaf phenotypes after the post-submergence phase: the old leaf was generally barely alive and severely damaged, whereas the young leaf fully recovered and continued to grow.

**Figure 2.**
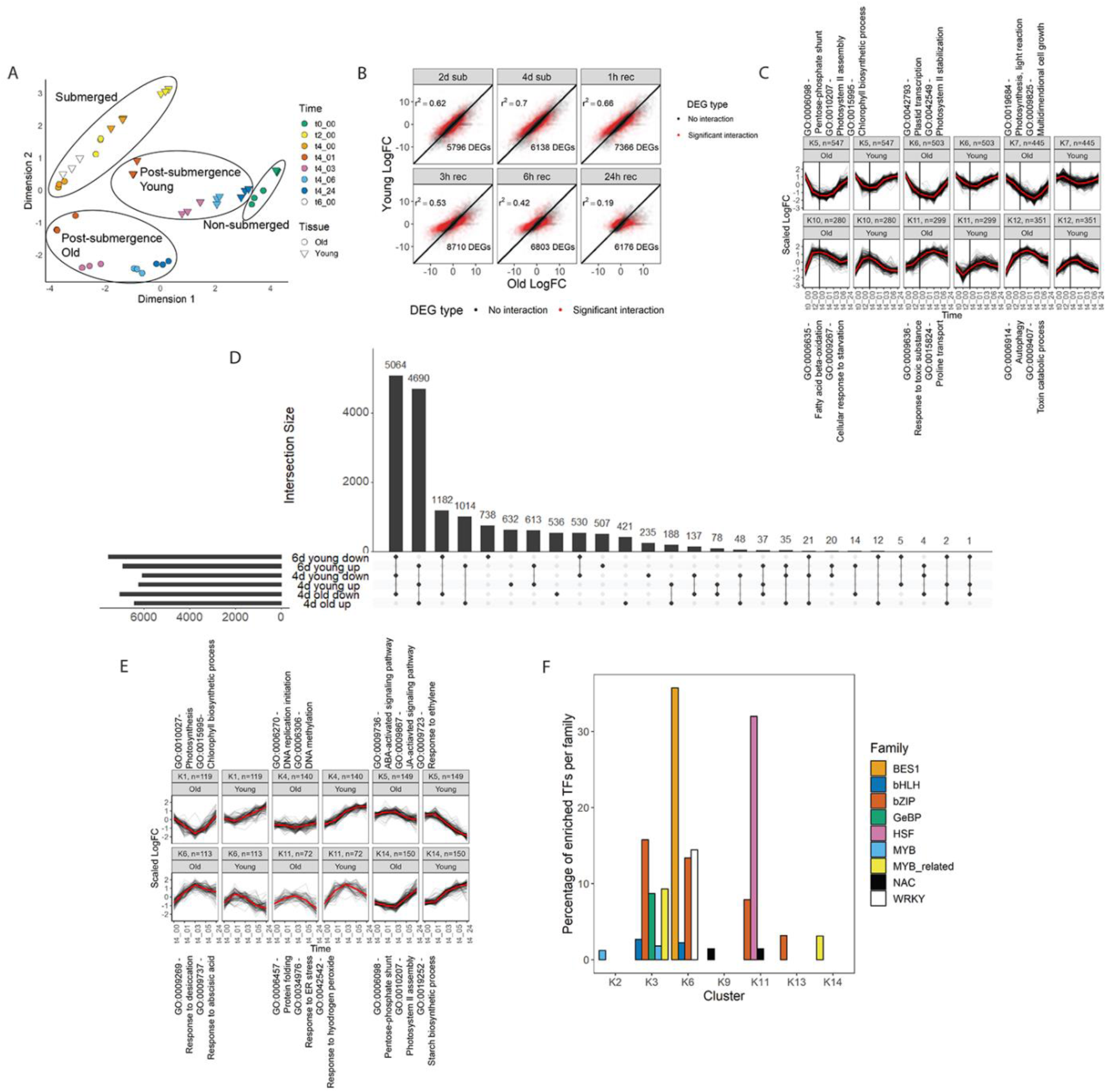
| Submergence and recovery both induce a leaf age-dependent reconfiguration of the transcriptome A) Multidimensional scaling (MDS) plot showing the distribution of the three biological replicates of the control (t0_00), submergence (t2_00, t4_00) and post submergence (t4_00, t4_01, t4_03, t4_06, t4_24) old and young leaf samples.B) Scatterplots comparing the response (log2FC) of individual genes in old and young leaves for the various submergence and recovery time points. Transcripts with a significant interaction for leaf age*time are highlighted in red, the number of these DEGs is shown in the bottom-right of each panel. Pearson’s r2 of the correlation between gene expression in old and young leaves in each timepoint is shown in the top-left of each figure. C) Clustering of the 4977 genes with significantly different expression between old and young leaves at a minimum of four timepoints across all sampled timepoints. Expression was calculated for old and young leaves relative to their respective expression under control conditions. A selection of overrepresented GO terms are shown per cluster. D) UpSet plot showing the intersection between genes induced or repressed after four or six days of submergence in young leaves and genes induced or repressed after four days of submergence in old leaves. Dots show which categories are intersected, numbers above each vertical bar indicate the size of each intersect. Horizontal bars indicate the size of each individual category. E) Clustering of the 1845 genes with significantly different expression between old and young leaves at a minimum of two timepoints across all post-submergence timepoints. Expression was calculated for old and young leaves relative to their respective expression at the last timepoint before desubmergence. A selection of overrepresented GO terms are shown per cluster. F) The percentage of transcription factors per family of which the targets are significantly enriched among the genes in clusters from Fig. 2E and Supplemental Fig. 6.

To analyze leaf age-dependent changes in gene expression, we also calculated DEGs between old and young leaves. At each of the six sampling timepoints during the submergence and post-submergence phases, more than 5000 genes showed a leaf age-dependent response to the treatment (Supplemental Fig. S1B). This number was highest at the three-hour post-submergence timepoint, where 8710 genes showed an age-dependent response. Although many genes responded in a leaf age-dependent manner to submergence, the correlation in gene expression between old and young leaves was relatively high during this phase (Fig. 2B). This correlation was largely lost during the post-submergence phase, indicating an age-dependent divergence of the transcriptomes during this recovery phase.

### Old and young leaves share some common responses to submergence and recovery

To explore the age-independent response to submergence, we first focused on the common stress responses between old and young leaves. We did this by selecting all genes whose expression was significantly different from t0_00 at two or more timepoints of the full time course (submergence and post-submergence), and selecting the genes in which the difference in response between old and young leaves was never significant (an FDR-adjusted p-value > 0.05 and/or an absolute log_2_ fold change < 1). This approach led to a selection of genes that change in expression during the submergence and post-submergence treatments, but in an age-independent way. The resulting list of 1629 genes was then divided over six clusters by fuzzy k-means clustering, which allowed us to identify common patterns in gene expression (Supplemental Fig. S2A). Genes with a maximum cluster membership lower than 0.50 were then omitted to reduce the noise in the dataset. The remaining 1482 genes, whose response to the treatments was independent from leaf age, were then further analyzed by testing which gene ontology (GO) terms are enriched per cluster (Supplemental Fig. S2B).

Several of the GO terms significantly enriched in cluster K1 were related to immune responses, like GO:0010200 “response to chitin” and GO:0002679 “respiratory burst involved in defense response”. Immune responses are known to be induced by submergence treatment (Hsu et al., 2013). This was consistent with the expression pattern of genes in cluster K1, which were upregulated gradually during submergence and slightly further upregulated during the post-submergence phase. Cluster K1 was also enriched for genes associated with endoplasmic reticulum stress, like those in categories GO:0034976 “response to endoplasmic reticulum stress” and GO:0006457 “protein folding”. Lastly, cluster K1 was enriched for genes associated with ethylene-related GO terms, like GO:0009723 “Response to ethylene”. The age-independent response to ethylene was likely a result of systemic accumulation of ethylene in submerged plants(Voesenek and Sasidharan, 2013; Sasidharan and Voesenek, 2015) .

Many of the GO terms enriched in cluster K2 were related to growth and photosynthesis, processes that are paused during submergence and reactivated during the post-submergence phase. This pattern of downregulation during submergence and reactivation during the post-submergence phase was also found in cluster K6, although the downregulation was much steeper here. Cluster K6 was enriched for GO terms involved in translation, consistent with previous reports that have found a rapid general shutdown of this process during low-oxygen stress although translation of specific transcripts is increased (Sachs et al., 1980; Juntawong et al., 2014; Cho et al., 2022). Clusters K2 and K6 were the two gene clusters with the most members (282 and 280 members, respectively), which highlights how common the shutdown of processes associated with growth and development was during submergence stress.

Surprisingly, we did not observe the enrichment of GO terms associated with hypoxia stress, such as GO:0001666 (“Response to hypoxia”). The expression of multiple core hypoxia genes varied significantly between old and young leaves, although there was no trend towards a stronger response in either leaf during submergence (Supplemental Fig. S3). During the post-submergence phase, however, expression of core hypoxia genes was reduced faster in young leaves than in old leaves (Supplemental Fig. S3).

### Age-dependent leaf responses to submergence and recovery

To identify the leaf age-dependent responses with a potential role in stress tolerance, genes that showed a significant age-dependent response to submergence or recovery at a minimum of four out of the six timepoints were selected. These were visualized by classification into 14 clusters by fuzzy k-means clustering, omitting the genes with a maximum cluster membership lower than 20% to reduce noise (Supplemental Fig. S4A, Fig. 2C). This revealed that the pausing of photosynthesis and growth during the submergence phase occurred more rapidly in old leaves than young leaves, as GO terms “Leaf morphogenesis”, “Plastid transcription”, and “Photosynthesis, light reaction” were overrepresented in clusters K5-K7 (Supplemental Fig. S4B, Fig. 2C). These processes were not only arrested faster in old leaves during the submergence phase, but they also resumed more slowly during the post-submergence phase. As expected, the induction of genes associated with senescence-related GO categories was stronger in old leaves than in young leaves (clusters K10 and K12) concomitant with the earlier initiation of senescence in older leaves (Rankenberg et al., 2024). Measurement for the enrichment of targets of transcription factors (TFs) per cluster revealed candidates regulating age-dependent responses. Of note was identification of clusters strongly enriched for targets of WRKY (clusters K2 and K13), and bZIP (Cluster K6) TFs (Supplemental Fig. S4C).

For all sampling timepoints (Fig. 1C) both an old leaf and a young leaf were harvested. The exception was the six days submergence timepoint, where only the young leaf was sampled since the old leaf was generally dead after six days of submergence. It was of interest to establish whether the transcriptome of a ‘near-death’ young leaf after six days of submergence was like that of a ‘near-death’ old leaf after four days of submergence. For this, the young leaf submergence effect DEGs after four days (6246 upregulated and 6094 downregulated genes) and six days (6934 upregulated and 7555 downregulated) of submergence were overlapped with the old leaf submergence effect DEGs after four days of submergence (6431 upregulated and 7053 downregulated). The majority of DEGs analyzed were significantly down- or upregulated in all three of these sets of genes (Fig. 2D). Of the genes that were not shared between all three sets (3652 for upregulated DEGs, 3635 for downregulated DEGs), most overlap was between the transcriptomes of old leaves of plants submerged for four days and young leaves of plants submerged for six days, with 1182 shared downregulated genes and 1014 shared upregulated genes, respectively. This was also consistent with the MDS plot in Fig. 2A, in which the transcriptomes of old leaves submerged for four days, and young leaves submerged for six days were located close to each other. Gene ontology analysis of these two sets of genes showed that processes commonly downregulated were often involved in fundamental cellular processes like transcription and translation (Supplemental Fig. S5). Shared upregulated genes between old and young leaves at their last submergence timepoint were involved in transport-related processes, suggesting export of the last nutrients out of the dying leaf for use in other parts of the plant. The high degree of similarity between the “near-death” transcriptomes of old and young leaves showed that a prolonged submergence treatment eventually had the same transcriptomic result in these leaves. The difference between the two leaves was in the time it took for them to reach this point.

### Age-dependent regulation of transcriptional responses during recovery

Clustering genes based on their expression patterns during the submergence and post-submergence phases allowed us to identify processes that are regulated in an leaf age-dependent manner. However, the dominant effect of the submergence phase might mask any age-dependent regulation that is specific to the post-submergence phase. To identify leaf age-dependent processes specific to the post-submergence phase, DEGs with a significant interaction effect for Time*Age at a minimum of three out of the four post-submergence timepoints were selected.

These 1848 genes were then divided over 14 clusters by fuzzy k-means clustering, and genes with a maximum cluster membership below 20% were again removed (Supplemental Fig. S6). Consistent with the results in Fig. 2C, the increase in transcripts of genes related to photosynthesis and growth was faster during recovery in young than in old leaves (Fig. 2E clusters K1, K4, and K14, Supplemental Fig. S6A and S6B). Cluster K5 showed that processes controlled by the stress-related hormones ethylene, ABA, JA, and SA were shut down faster in young leaves than old leaves during the post-submergence phase.

Clustering of the age-dependent responses to the recovery phase revealed a stronger and more sustained induction of genes associated with the GO categories “Response to desiccation” and “response to abscisic acid” in old leaves (Fig. 2E, cluster K6). This was consistent with the visual phenotype of plants recovering from submergence, where there was more desiccation in old than in young leaves (Fig. 1A). ABA-mediated responses to dehydration are controlled by ABA-RESPONSIVE ELEMENT BINDING FACTOR (ABF) TFs of the bZIP family (Uno et al., 2000). The targets of several ABFs, and other bZIP TFs, were overrepresented among the genes in cluster K6 (Fig. 2F).

Cluster K11 showed a strong enrichment for multiple GO categories associated with protein misfolding and exposure to hydrogen peroxide, two processes that are often connected (Ozgur et al., 2018). Expression of genes in this cluster increased further in young leaves than in old leaves during the post-submergence phase. Targets of HEAT SHOCK FACTOR (HSF) and bZIP TFs were overrepresented in cluster K11 of genes responding to the post-submergence phase, which was consistent with the involvement of TFs of these families in these processes (Fig. 2F, (Jung et al., 2013; Ko and Brandizzi, 2022). Of the HSF TFs whose targets are overrepresented in cluster K11, some had clear leaf age-dependent patterns (Supplemental Fig. S7). The expression of both *HSFA6A* and *HSFA6B* was higher in young leaves than in old leaves during the first hours post-submergence. These two closely related TFs play a role in the interaction between ABA signaling and ROS signaling (Wenjing et al., 2020). Their higher expression in young leaves could facilitate their faster response to desubmergence, as ABA and ROS signaling are both important during submergence recovery (Yeung et al., 2019). In maize, the expression of a homolog of Arabidopsis *HSFA6B*, *HSFTF13* is induced by bZIP60 during heat stress (Li et al., 2020). Under heat stress, HSFTF13 induces the expression of HEAT SHOCK PROTEINS (HSPs), which function as chaperones for accumulated misfolded proteins under stress conditions. This is also consistent with the overrepresentation of genes associated with ER stress and protein folding in cluster K11 (Fig. 2E; Supplemental Fig. S6).

### Age dependent leaf dehydration during recovery can be attributed to differential ABA sensitivity and leaf conductance

During recovery, older leaves visibly dehydrated faster than young leaves (Fig. 1A). This trend was confirmed from fresh weight and water content measurements of old and young leaves recovering from different submergence durations (Fig. 3A, 3B). These two parameters of leaf dehydration thus confirmed more frequent dehydration in old leaves than in young leaves (p<0.05, Fisher’s exact test). There was a positive correlation between dehydration and submergence duration. This was likely due to the inability of submergence damaged leaves to recover post-submergence. Consistent with this age-dependent leaf dehydration during the post-submergence phase, several genes associated with this process were regulated in an age-dependent manner (Fig. 2E). Notably, Cluster K6 with sustained upregulation in old leaves and transient induction young leaves (Fig. 2E), was enriched for genes associated with the GO category “response to ABA” (GO:0009737). To further evaluate this trend in correlation with differential leaf dehydration during recovery, we assessed the expression of 250 ABA responsive genes(Cao et al., 2017) during recovery in our dataset (Fig. 3C). Counterintuitively, we found an inverse correlation between ABA transcriptional response and dehydration. While old leaves showed a consistent ABA response across all recovery time points, the corresponding response in young leaves was transient. An assessment of the expression of ABA metabolic genes during recovery suggested no major leaf age-dependent differences in ABA metabolism during recovery (Fig. 3D). As degradation of the cuticle during submergence has been associated with post-submergence water loss (Xie et al., 2020), we investigated whether submergence affected the permeability of the leaf cuticle in an age-dependent manner. Using toluidine blue uptake as a proxy (Tanaka et al., 2004), we observed a similar increase in cuticular permeability in both old and young leaves (Supplemental Fig. S8).

**Figure 3.**
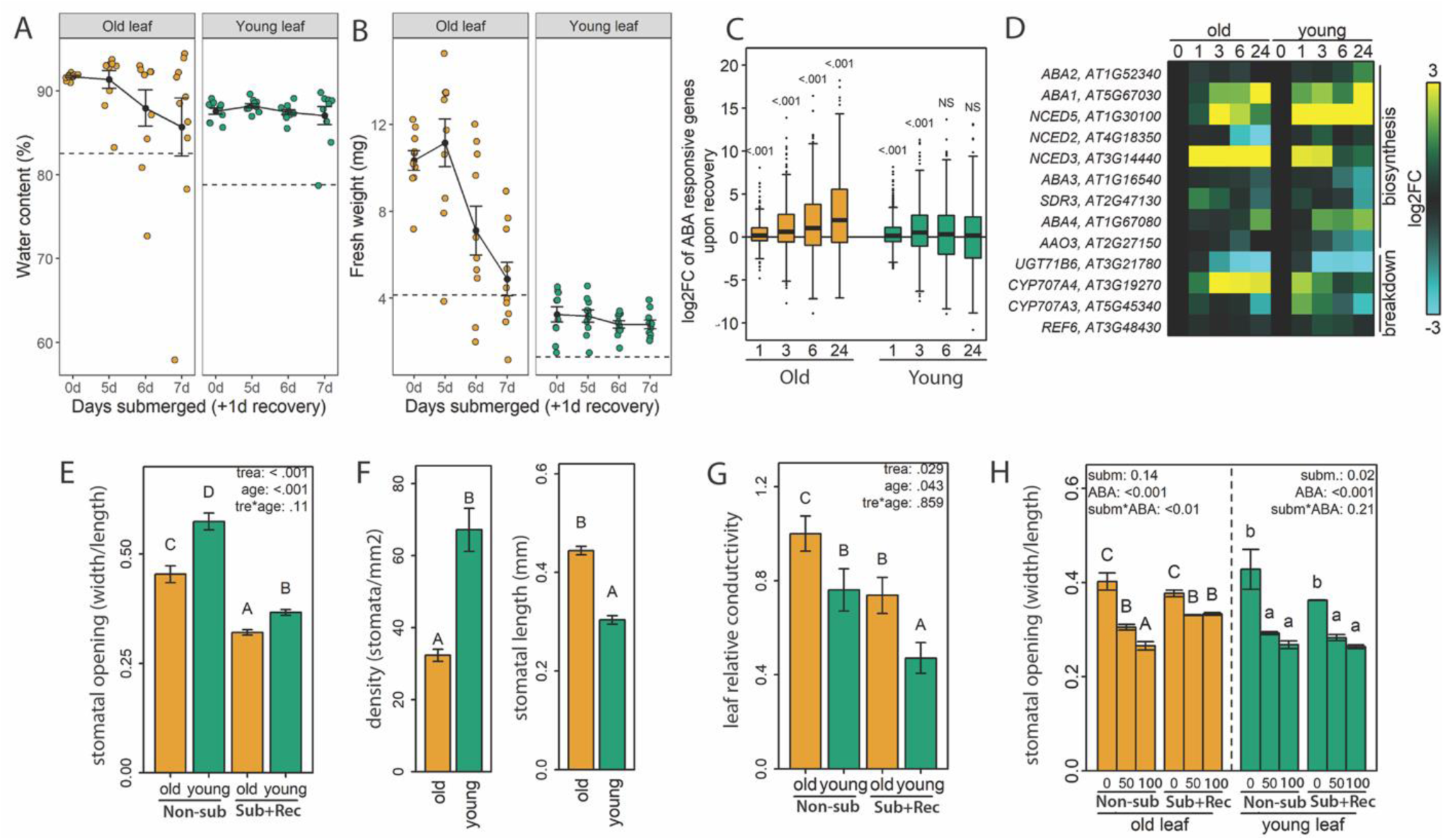
**| Differential ABA sensitivity and leaf conductivity contribute to age-dependent dehydration during post-submergence recovery** A) Water content (the percentage of water per leaf weight) of old and young leaves one day after the indicated submergence durations. The dashed line indicates 90% of the mean water content of leaves of non-submerged plants, leaves with lower water content were designated as dehydrated. This happened more often in old leaves than in young leaves (p<0.05, Fisher’s exact test), n = 10 leaves. B) Fresh weight of old and young leaves one day after the indicated submergence durations. The dashed line indicates 40% of the fresh weight of leaves under control conditions, leaves with a lower fresh weight were designated as dehydrated. This happened more often in old leaves than in young leaves (p<0.05, Fisher’s exact test), n = 10 leaves. C) The response to recovery of genes considered ABA induced. Genes that were among the top 250 induced genes 6 hours after, and top 250 genes 24 hours after 24 ABA treatment in leaf tissue (Cao et al. 2017) were considered ABA induced (441 genes). Significance values above boxplots concern, for the corresponding timepoint, the enrichment of ABA induced genes among recovery induced genes compared to non-recovery induced genes (Fisher’s exact test). Timepoints shown are hours post-submergence recovery. D) Heat map showing the expression of ABA biosynthesis and degradation genes at various timepoints following desubmergence (0, 1, 3, 6 and 24h). E) Stomatal aperture of old and young leaves before and after four days of submergence followed by three hours of recovery. N= 166-192 stomata per sample F) Stomatal length and stomatal density on the adaxial side (n= 18-25 images per leaf type) of old and young leaves. G) Relative difference in leaf conductivity between experimental groups. Here the relative conductivity is based exclusively on water loss via stomates. This amounted to the combined effect of the density of stomates, the area of each pore and Poiseuille law that stated that laminar flow rate scales quadratic to the area. Pore area was estimated by an ellipse calculated from the stomatal pore width and length. Calculations are based on data presented in E and F. H) Stomatal aperture of old and young leaves before and after four days of submergence after treatment with 0uM (mock), 50 uM or 100 uM ABA (for 3h). Per sample, between 350-700 stomata were measured from 3-5 leaves per group.

For **E-H** the data are mean ± sem (n=5). Significance of the two-way ANOVA effects are given in top corner of the graphs. Here “trea” is treatment effect, “non-sub” the control non-submerged treatment and and sub+rec as submergence followed 3h of recovery. Age is the effect of either a young or old leaf. ABA is the effect of mock (0 uM) or ABA treatment (50 or 100 uM). Tre*age and subm*ABA indicate the corresponding interaction effects. Letters indicate differences between individual bars (p <0.05).

Next we probed the possibility that faster dehydration of old leaves might be explained by a larger stomatal aperture or higher stomatal density. An assessment of stomatal traits for young and old leaves revealed that while stomatal aperture changes were similar for old and young leaves after 3 hours of recovery (Fig. 3E), the latter had smaller stomatal size but higher stomatal density (Fig. 3F). Leaf conductance is the net effect of pore size and density, where individual pore area scales quadratically with conductance (Poiseuille law). The combination of density, pore size and pore aperture indicated that old leaves would indeed have a higher conductance during the submergence recovery phase (Fig. 3G).

We also tested whether dehydration differences between old and young leaves could be attributed to differences in ABA sensitivity. We found that ABA sensitivity of old leaves was significantly reduced during the post-submergence phase (Fig. 3H). Whereas a submergence treatment did not affect the closure of guard cells in response to ABA in young leaves, it reduced the effect of ABA on the stomatal aperture of old leaves. Thus in conclusion, the observed faster dehydration of old leaves is likely a combination of higher conductance and reduced ABA sensitivity following submergence.

### ER stress signaling contributes to recovery of young leaves

At the submergence durations used here, old leaves dehydrate, are unable to recover, and die. In contrast, younger shoot tissues resume growth following desubmergence. Among the biological processes identified as differentially regulated in young leaves were genes associated with ER stress and protein folding (Fig. 2E, Fig. 4A). We hypothesized that ER stress sensing and mitigation via activation of the unfolded protein response (UPR) during and after submergence contribute to post-submergence recovery in younger tissues.

**Figure 4.**
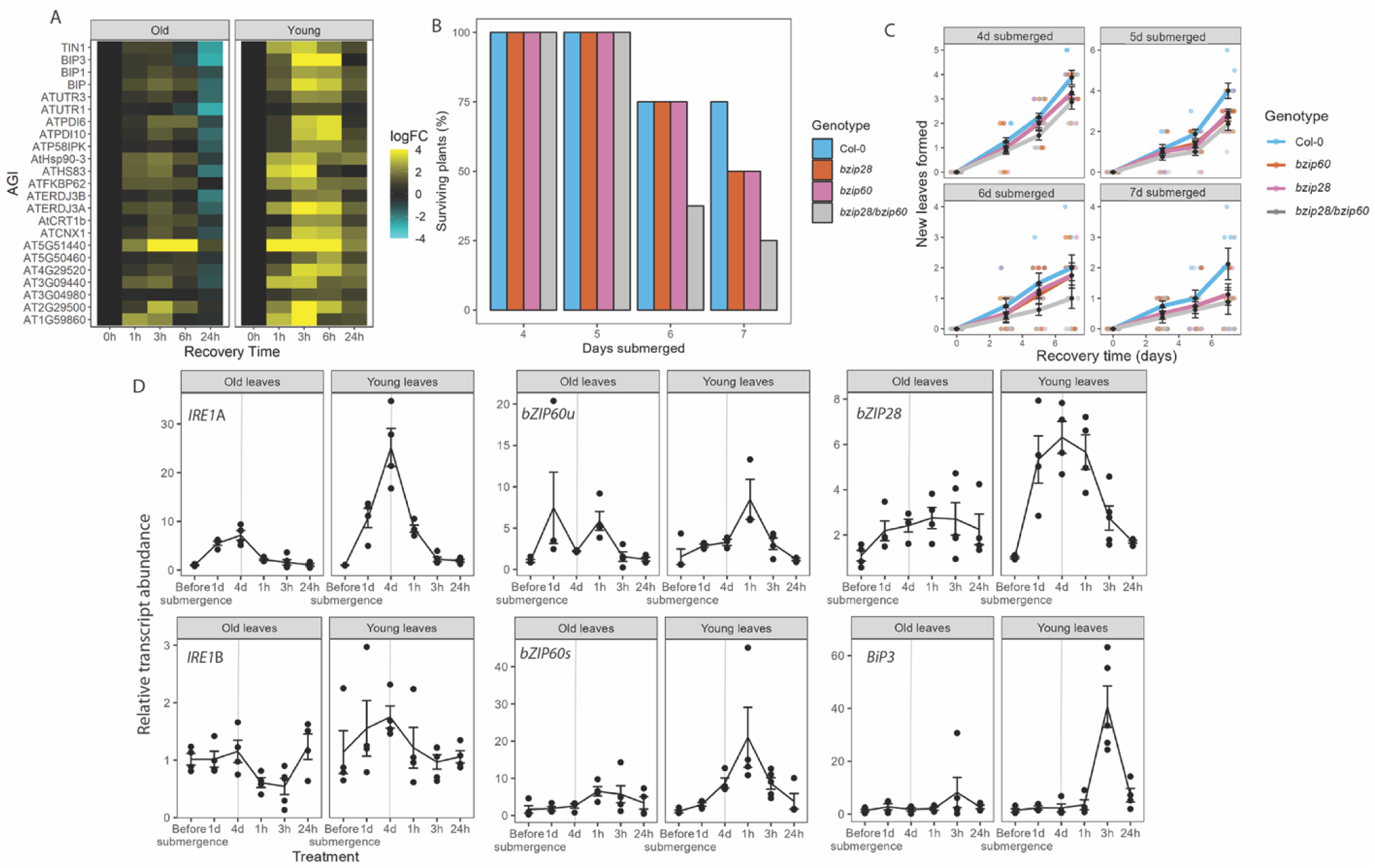
**| ER stress signaling contributes to post-submergence recovery in young tissues** A) Expression of transcripts in cluster 11 (Fig. 2E, Supplementary Fig. S6) associated with protein folding (GO:0006457) in young and old leaves during recovery, plotted relative to respective expression at various recovery durations following four days of submergence. B) Survival of Col-0, *bzip28*, *bzip60*, and *bzip28/60* plants after different submergence durations. Plants were submerged at the 10 leaf stage, desubmerged at the indicated submergence duration and allowed to recover under control growth conditions for seven days. At this point survival was scored, with plants being scored as dead or alive based on the absence or presence of visible green tissues. At each time point survival was scored for eight plants.C) New leaf formation of Col-0, *bzip28*, *bzip60*, and *bzip28/60* during recovery from different submergence durations. A significant genotype effect was observed during recovery, as determined by a two-way ANOVA for recovery time* genotype. Per time-point, new leaf formation was scored for n= 8 plants. E) Transcript abundance of ER stress signaling pathway genes on old and young leaves during submergence (1d, 4d) and recovery following 4d of submergence (1h, 3h, 24h). Gray vertical line indicates end of submergence period and start of recovery. Mean is average of 3 biological replicates (n= 3) with each consisting of 5 young or 5 old leaves from a single plant.

The ER is the primary site of oxidative protein folding, achieved via a disulfide relay system comprising protein disulfide isomerase (PDI) and ER oxidoreductin (ERO) enzymes. As molecular oxygen is the terminal electron acceptor for EROs, disulfide bond formation is sensitive to hypoxia (Ugalde et al., 2022). ER stress is triggered by stresses such as submergence and the concomitant reduction in oxygen availability (Gayral et al., 2017; Zhou et al., 2021; Gibbs et al., 2024) leading to an accumulation of misfolded or unfolded proteins in the ER. This elicits the UPR, which serves to restore homeostasis. Two classes of ER stress sensors are known in plants: membrane associated basic leucine zipper (bZIP) TFs and RNA splicing factors, specifically Inositol-requiring enzyme type 1 (IRE1). We tested the physiological significance of the observed ER-stress markers inductions using single and double mutant combinations for *bZIP60* and *bZIP28*, each mediating the two UPR activating pathways (Ko and Brandizzi, 2022). bZIP28 is an ER stress sensor that directly mediates nuclear activation of stress responsive genes. bZIP60 operates downstream of IRE1, activating the UPR following a splicing event that permits its translocation to the nucleus. We observed no developmental phenotypes associated with these mutations. However, there was a clear negative effect of submergence correlated with stress duration (Fig. 4B, Fig. 4C). Consistently, plant survival of the double mutant was reduced by more than 50% when plants were submerged for more than 6 days (Fig. 4B). Despite these phenotypes, all genotypes recovered and were able to produce new leaves (Fig. 4C). Nevertheless, this capacity for shoot regeneration was significantly diminished in *bzip28/bzip60* especially after long-term submergence (Fig. 4C). These results indicated that having two functional branches of the UPR is required for shoot regeneration and plant survival after de-submergence. Notably, the degree of senescence was not affected since there were no significant differences in chlorophyll content between the genotypes at any time point (Supplemental Fig. S9).

To test the involvement of ER stress responses at the molecular level, we measured levels of ER-stress markers in old and young shoot tissues by qRT-PCR (Fig. 4D). Time-course analysis showed a more prevalent accumulation in younger leaves than in older tissues. Moreover, rapid accumulation of ER-stress sensors *IRE1a*, *IRE1b*, and *bZIP28* was evident within 1d of submergence and maximal by the end of treatments (4d). Levels of *bZIP60*u, a splicing target of IRE1 were greatest 1h after de-submergence, as were the levels of the spliced version (*bZIP60*s). Lastly, *BiP3*, a UPR gene induced by bZIP28 and bZIP60 was distinctly expressed only in the recovery period, 3h after de-submergence. Similarly, mRNAs for other UPR genes were distinctly enhanced only after de-submergence (Supplemental Fig. S10). This indicated that mRNAs for ER-stress sensing and UPR accumulate at distinct phases of submergence. ER-stress signals initiate during submergence while the UPR is induced only during the recovery phase in young shoot tissues.

### The activation of ER stress signaling in young leaves is specific to the recovery phase

To further dissect the contributions of ER-stress signals to the submergence response of Arabidopsis, we quantified whole genome and proteome effects. We hypothesized that transcriptome and proteome reconfiguration during submergence and recovery involves a combination of ER-dependent and -independent signals that are distinct in old and younger leaves. To determine the onset of responses that require ER stress signaling, we harvested old and young halves of Col-0 and *bzip28/60* rosettes before, during, and after submergence (Fig. 5A). Differential responses between each treatment and tissue type were separated over time by multidimensional scaling (MDS), whereas the genotype-dependent responses were smaller (Fig. 5B; Supplemental Table S2). Genotype-dependent responses were determined at each tissue and timepoint, and genes with an FDR-corrected p-value below 0.05 and an absolute fold change between the two genotypes were deemed as significantly different (Fig. 5C). Among the genes different between Col-0 and *bzip28/60*, genes in the GO category “Response to endoplasmic reticulum stress” (GO:0034976) were enriched in the set with a higher expression in Col-0 old leaves after four days of submergence, and after three hours of recovery in both old and young leaves. This shows that the response to ER stress is gradually activated in old leaves during submergence and post-submergence recovery, whereas the response is only induced in young leaves during the post-submergence phase. Consistent with this, among genes that are significantly co-expressed with the ER stress marker gene *BIP3* are several that are known to play a role in ER stress signaling (Fig. 5D). These all follow the pattern of a gradual induction during submergence that continues during recovery in old leaves, and a recovery-specific induction in young leaves.

**Figure 5.**
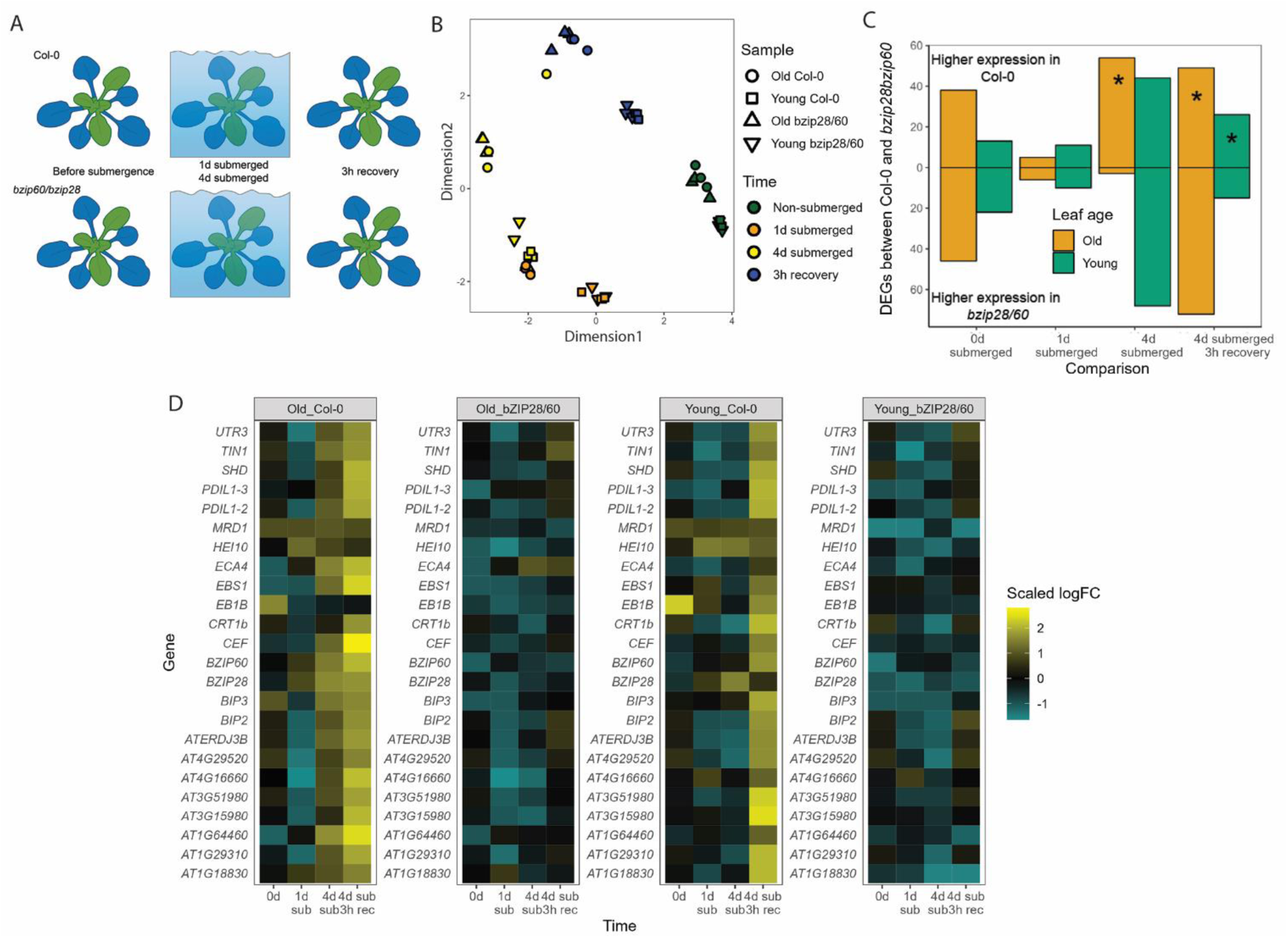
| **The role of ER stress signaling in transcriptional responses to submergence and recovery** A) Schematic of the harvested materials, young tissues are highlighted in green, old tissues are highlighted in blue B) MDS plot of the old and young transcriptomes of Col-0 and *bzip28/60* before, during, and after submergence. C) The number of genes differentially expressed between Col-0 and *bzip28/60* in old and young leaves at each timepoint. Columns highlighted with an asterisk are significantly (p<0.001) enriched for genes associated with the GO term “Response to endoplasmic reticulum stress” (GO:0034976) D) Genes of which the expression pattern is significantly (p<0.05) correlated with that of the ER stress marker gene *BIP3*, as determined by Pearson’s correlation coefficient. These contain multiple genes previously identified as being involved in the ER stress response, like *BIP2*, *UTR3*, *ATERDJ3B*, and *CTR1b*.

### Impact of the UPR on proteome responses to submergence-recovery stress

After establishing the consequences of the misregulation of ER stress signaling on the transcriptome, we sought to determine the global effects on the proteome. Considering that the demand for protein synthesis and oxidative protein folding in the ER is likely higher in young leaves and since the *bzip28/60* double mutant showed a slower rate of post-submergence leaf formation (Fig. 4D), we focused on the changes in the proteome of young leaves. We conducted two experiments using isobaric Tandem Mass Tag (TMTpro™) labelling to identify proteins with altered abundance in wild type and mutant in response to submergence-recovery treatments. Firstly, young leaves were harvested before and after one day of submergence. Dimensionality reduction via MDS showed a clear separation of samples by treatment and genotype (Fig. 6A). 3,608 proteins represented by a minimum of two peptides were identified and quantified (Supplemental Table S3; Supplemental Fig. S12A). The proteome after one day of submergence was largely similar in Col-0 and *bzip28/60*, but with lower fold-changes in the mutant (Fig. 6B, 6C, 6D). Relatively few proteins increased in abundance, as defined by a fold change (FC)>2 at p<0.05; 49 in Col-0 and 46 in *bzip28/60* (Fig. 6C and 6D). Isobaric labelling datasets tend to under-estimate fold-changes (Rauniyar and Yates, 2014) and applying a more relaxed cut-off (FC>1.5; p<0.05) identified 118 and 89 proteins with increased abundance following submergence in wild type and mutant, respectively (Supplemental Fig. S13A). In Col-0, these included proteins associated with sugar starvation responses: PER21, STY46, MCCA, IVD, KING1, TPS8, UGE1, BXL1, BGAL4, PCAP1, DRM1, ASN1 (Arias et al., 2014; Tarancón et al., 2017) and two ethylene biosynthetic enzymes, ACC oxidase (ACO)1 and ACO2. Only three of the Core 49 hypoxia proteins were up-regulated under submergence (Mustroph et al., 2009) but interestingly, five proteins regulated by PRT6, a candidate E3 ligase belonging to the oxygen-sensing N-degron pathway (Zhang et al., 2015) increased in response to submergence. A greater number of down-regulated proteins was identified: 120 in Col-0 and 55 in *bzip28/60*; FC>2; p<0.05; 316 and 208 at FC>1.5; p<0.05. Col-0 proteins with decreased abundance upon submergence were enriched in KEGG pathways associated with photosynthesis, flavonoid biosynthesis, and N-glycan biosynthesis and protein processing in the ER. The latter included Calnexin (CNX)1 and CNX2, a SecY family protein and members of the oligosaccharyl transferase complex. Mitochondrial membrane proteins [including voltage dependent anion channels (VDACs), TOM40-1, TOM9-2, MPT3 and DTC], and proteins associated with chromatin binding (including nucleosome assembly proteins, subunits of the FACT complex and ISWI chromatin-remodeling complex ATPase) were also down-regulated under submergence.

**Figure 6.**
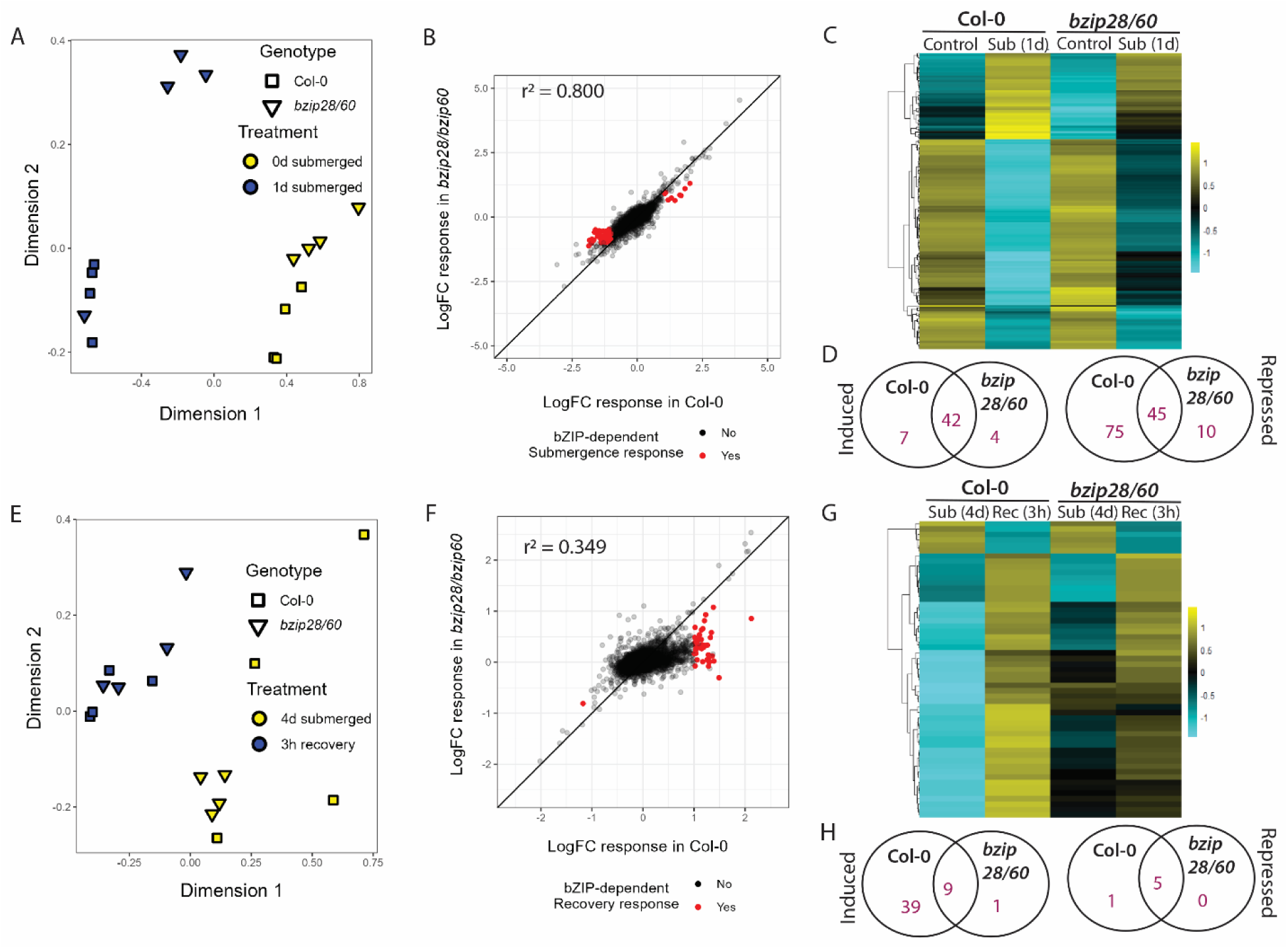
| **The role of ER stress signaling in submergence and post-submergence proteostasis** (A) MDS plot of the proteomes of Col-0 and *bzip28/60* before and after one day of submergence; proteins with at least 2 unique peptides are shown. (B) The response of proteins to one day of submergence in Col-0 and *bzip28/60*. Proteins that show a response in Col-0 but not in *bzip28/60* are highlighted in red. Pearson’s r^2^ of the correlation in protein LogFC in both genotypes is shown in the top-left corner. (C) Heatmap of scaled protein abundance for 183 proteins with at least 2-fold changes (p < 0.05) in one comparison of Col-0 and *bzip28/60* before and after one day of submergence D) Overlap between proteins induced and repressed in both genotypes after one day of submergence. (E) MDS plot of the proteomes of Col-0 and *bzip28/60* after four days of submergence and three hours of recovery. (F) The response of proteins to three hours of post-submergence recovery in Col-0 and *bzip28/60*. Proteins that show a response in Col-0 but not in bzip28/60 are highlighted in red. Pearson’s r^2^ of the correlation in protein LogFC in both genotypes is shown in the top-left corner. (G) Heatmap of scaled protein abundance for 55 proteins with at least 2-fold changes (p < 0.05) in one comparison between Col-0 and *bzip28/60* after four-day submergence and three hours of post-submergence recovery. H) Overlap between proteins induced and repressed in both genotypes after three hours of post-submergence recovery.

As genes involved in ER stress signaling were mostly induced during the post-submergence recovery phase (Fig. 5D), we also mapped the changes in the young leaf proteome during the first three hours of post-submergence recovery after 4 days of submergence. 4,767 proteins represented by a minimum of two peptides were identified and quantified (Supplemental Table S4; Supplemental Fig. S12B) and again, the samples grouped together based on both genotype and treatment (Fig. 6E). The proteome after 3h recovery was largely similar in Col-0 and *bzip28/60*, but with more deviation than the submergence response and lower fold-changes in the mutant (Fig. 6F, 6G, 6H; Supplemental Fig. S13B). In the wild type, many of the protein categories that were downregulated under submergence increased in abundance after recovery, most notably mitochondrial membrane proteins (including VDACs), photosynthetic proteins, proteins associated with chromatin and splicing, and protein processing in the ER (Supplemental Fig. S14). Whilst the data did not bear the strong hallmark of the classical UPR response seen in the transcriptome, the proteome of the *bzip28/60* mutant had not recovered to the same extent as wild type after 3h (Fig. 6F, Fig. 6G; Supplemental Fig. S13B), consistent with a defect in ER protein processing. Relatively few proteins were downregulated upon recovery, although abundance of three photoreceptors: cryptochrome1, phytochrome A and phototropin-2 was reduced in both mutant and wild type. Comparing scaled protein abundances revealed that VDAC1, 2, 3 and 4 are down regulated under submergence and accumulate during recovery, an effect that is less pronounced in the *bzip28/60* mutant compared to wild type (Supplementary Fig. S15A). This prompted us to investigate the impact of VDAC function on submergence-recovery resilience. Rosettes at the 10-leaf stage were subjected to 6 d submergence and new leaf emergence was monitored over 3, 7 and 10 d of recovery (Supplementary Fig. S15B-C). As already demonstrated for wild type plants (Fig. 1;(Rankenberg et al., 2024)), *vdac1* and *vdac2* older leaves died and younger leaves and the meristem remained green. Monitoring new leaf formation over 10 d of recovery revealed that compared to wild-type Col-0 plants, *vdac1* was impaired in new leaf formation following submergence, whereas *vdac2* and *vdac3-1* mutants had a modest but statistically significantly increased rate of leaf formation (Supplementary Fig. S15; Supplemental Table S5). Taken together, these data suggest that VDACs play distinct but overlapping roles in recovery from submergence.

## Discussion

Plant age, tissue and cell-type identity can be important determinants of response and tolerance to environmental stresses. This is also the case for plant responses to flooding. Previous studies have explored the specific responses of plant organs or cell types to submergence and hypoxia, but few have addressed the effect of leaf age (Mustroph et al., 2006; Van Veen et al., 2016; Müller et al., 2021; Geldhof et al., 2024; Rankenberg et al., 2024) . In this study we sought to characterise the molecular processes mediating the leaf-age-dependent stress responses of *Arabidopsis thaliana* during submergence and recovery. The Arabidopsis rosette offers an excellent system to probe leaf-age dependent responses to submergence. Upon complete submergence, differential stress symptoms are clearly visible in terms of senescence and dehydration. During recovery, progression of symptoms occurs only in old leaves while young leaves recover growth and meristem activity is restored. Exploiting these differences, we sought to understand the molecular bases for this stress phenotype.

### Leaf-age dependent responses are most evident during recovery corresponding with significant transcriptomic modulation

During complete submergence, in theory, the stress is imposed equally across the entire rosette. For example, initial changes related to submergence such as mechanical stress from the water column or the accumulation of ethylene are expected to be equal across the rosette. The assessment of the Arabidopsis rosette transcriptomes revealed substantial remodelling under stress. Surprisingly, young and old leaf responses showed a significant correlation during submergence (Figure 2C). These core responses included several molecular processes typically activated in response to submergence stress including response to ethylene and the downregulation of growth, photosynthesis and translation. An exception was the response to hypoxia which showed strong leaf age-dependent responses, although no clear trend was observed (Supplemental Fig. S3). The similarity between the transcriptomes of a 4d-submerged old leaf and a 6d-submerged young leaf suggested that the difference between old and young leaves in their response to submergence was in part determined by the speed of their response. Submergence treatment thus induced “transcriptomic aging” of a leaf and old leaves were more sensitive to this induction than young leaves (Fig. 2D).

Leaf-age dependent transcriptome responses were most apparent during the recovery phase. The recovery period following submergence can be viewed as a sequential stress for plants. Transition to aerial light and oxygen conditions is associated with increased dehydration and oxidative stress (Yeung et al., 2018; Yeung et al., 2019). Young leaf transcriptomes were in general marked by a better recovery (Fig. 2A) suggesting a higher tolerance of post-submergence stress and in keeping with the observed phenotypes. Analyses of the responses specific to the post-submergence phase confirmed the induction of several pathways previously implicated in recovery. Quantitative differences in post-submergence dehydration and ROS accumulation contribute to differential submergence tolerance of the Arabidopsis accessions Bay-0 and Lp2-6 (Yeung et al., 2018). In the current study, the response to these stimuli was induced more strongly in young leaves than old leaves (Fig. 2E, Supplemental Fig. S4). In addition, young leaves showed a stronger induction of genes associated with ER stress and the UPR. While old and young leaves both repressed processes associated with growth and photosynthesis during submergence, young leaves were able to restart these processes faster upon desubmergence. Many TFs were regulated in an age-dependent manner although there were surprisingly few that exhibited opposite expression patterns between old and young leaves. This also applied to non-TF genes: differences between old and young leaves were mostly in the strength of the response rather than the direction of the response.

### Senescence and dehydration are initiated in older leaves

The ability to delay foliar senescence (the “stay-green” phenotype) is considered a valuable trait for sustained yield and biomass under control and stress conditions (Kamal et al., 2019), and accordingly, the mechanisms underlying age-dependent leaf senescence have been extensively studied (Schippers, 2015). Submergence presents a unique case where despite systemic accumulation of ethylene (a well-known positive regulator of senescence) chlorophyll degradation still occurs in a leaf-age dependent manner. Mechanisms underlying this leaf age-dependent response to a leaf age-independent signal are not fully resolved (Rankenberg et al., 2021; Rankenberg et al., 2024). Consistent with the observed phenotype, processes related to senescence were induced faster in old leaves, but they were not enriched among genes with higher expression in old leaves under control conditions (Supplemental Fig. S4). This suggests that the earlier onset of submergence-induced senescence in old leaves was not merely due to baseline differences in development but caused by differences in the speed at which senescence was activated. Age-related differences in the onset of senescence are associated with age-related changes (ARCs) that control when a leaf is sensitive to senescence-inducing signals(Jibran et al., 2013; Schippers, 2015; Kanojia et al., 2020). An important regulator of chlorophyll catabolism in Arabidopsis is the NAC TF ORE1 (Qiu et al., 2015; Rankenberg et al., 2024). Although *ORE1* expression was strongly induced in both old and young leaves during submergence, its expression declined during the post-submergence phase in young leaves only (Supplemental Fig. S16). The gradual derepression of ORE1 by *miR164* with leaf age is a prime example of these ARCs (Jin et al., 2009) but few other mechanisms have been identified. We showed recently that post-translational ORE1 regulation mediates differential leaf-ageing during submergence, in a process that does not involve *miR164* (Rankenberg et al. 2024).

In the post-submergence phase, accelerated senescence can be linked to increased dehydration. Plants recovering from submergence stress typically experience physiological drought, as their damaged root systems are unable to compensate for water lost via the leaves (Yeung et al., 2019; Liu and Zwiazek, 2022). Consistent with the observed phenotype, older leaves of submerged plants had an unfavourable water status compared to young leaves and stronger enrichment of clusters related to dehydration (Fig. 3A). Flooding is known to induce molecular and physiological responses similar to drought in various species, including a strong upregulation of ABA in shoots during waterlogging and submergence recovery (Yeung et al., 2018; Zagorščak et al., 2025). However, the exact role of ABA in during submergence recovery and the correlation with drought tolerance remains unclear. While our transcriptome data revealed a more persistent ABA response in old leaves during recovery, there were no indications for differential transcriptional regulation of ABA biosynthesis. Greater post-submergence dehydration in the flood sensitive Arabidopsis accession Bay-0 correlated with higher ABA levels and ABA-mediated induction of the guard cell localised *SAG113* (Yeung et al., 2018). SAG113 mediates greater stomatal opening and water loss in older leaves to accelerate senescence. The higher expression of *SAG113* in old leaves in our dataset pointed to a similar mechanism underlying the greater dehydration of older leaves during recovery (Supplemental Fig. S16). However, rather than stomatal movement and density, our data suggests that leaf-age dependent dehydration differences during recovery are explained by a combination of submergence-mediated reduction of ABA sensitivity and higher leaf conductance in old leaves.

### ER stress responses contribute to submergence recovery in young leaves

During adverse conditions, the proper folding of secretory proteins in the ER can be disrupted, leading to an overload of misfolded proteins in the ER lumen and causing ER stress. This condition is managed by initiating the UPR which serves to maintain ER homeostasis by aiding removal of misfolded proteins and enhancing protein folding capacity. A swift response to ER stress is essential for restoring cellular homeostasis and stress recovery. If the response is inadequate it can culminate in programmed cell death. UPR signaling in plants is mediated by two routes: involving the ER membrane-associated TFs bZIP17 and bZIP28 and the ER membrane localized RNA-splicing factors IRE1/2 (Ko and Brandizzi 2024). Considering the oxygen-dependency of protein folding in the ER (Fuchs et al., 2022), it is not surprising that flooded tissues experience ER stress. The induction of ER stress-associated transcripts observed here was consistent with previous studies showing the importance of the UPR for hypoxia and flooding (Zhou et al., 2021; Ugalde et al., 2022) . However, our data suggests this has an underlying age-dependency.

Our observation of a stronger ER stress signature in young leaves during recovery could suggest more severe ER stress (perhaps related to higher protein synthesis rates compared to old leaves) or a greater capacity to respond to ER stress. Assessment of mutants disabled in the two branches of the UPR signaling pathway revealed an impact on survival following long-term submergence characterized by a difference in new leaf formation, but not in senescence (which occurs in old leaves). This suggests that while flooding likely induced accumulation of misfolded proteins leading to ER stress in both old and young leaves, young tissues demonstrate ER stress recovery and adaptive strategies to permit continuation of growth, as reflected in resumption of new leaf formation. This was also supported by a stronger induction of upstream regulators (*IRE1a*, *bZIP60s*, *bZIP28*) and their downstream targets (e.g. *BiP3*) during recovery in young leaves. An assessment of the impact of ER stress on the proteome of young leaves revealed that protein processing in the ER is down regulated upon submergence in agreement with the oxygen-dependency of ER protein folding. However, initiation of UPR pathways as plant tissues transition to recovery conditions is essential as evidenced by a slower recovery in *bzip28/60* because the UPR is impaired. Our data also indicate that while ER stress is likely sensed during submergence, UPR activation occurs during recovery. While not investigated here, this could be associated with the generation of signals such as ROS during this phase (Yeung et al., 2018; Gibbs et al., 2024; Khan et al., 2024; Yu et al., 2024).

The proteome comparisons between wild-type and UPR mutants also pointed to a role for mitochondrial proteins during submergence recovery. Mitochondrial function is dramatically modulated during submergence-recovery. Reduced oxygen availability under submergence inhibits the mitochondrial electron transport chain and subsequently, the restored respiratory capacity upon reoxygenation leads to a burst of ROS production (Chang et al., 2012; Sasidharan et al., 2018; Khan et al., 2024). Amongst the mitochondrial proteins whose abundance was affected both by the stress and the disruption of ER stress signaling were the VDACs. VDACs are the most abundant proteins of the mitochondrial outer membrane (Fuchs et al., 2020) and have overlapping and distinct functions in mitochondria (Tateda et al., 2011; Robert et al., 2012; Hemono et al., 2020) not only being involved in exchange of metabolites and ions between the cytosol and the intermembrane space, but also playing roles in tRNA import and oxidative stress responses (Salinas et al., 2006; Hemono et al., 2020; Kanwar et al., 2022). The Arabidopsis genome encodes six VDAC genes, of which only four are expressed (Tateda et al., 2011; Robert et al., 2012). Assessment of the role of the VDACs revealed a role for VDAC1, the most abundant VDAC in the resumption of new leaf formation following desubmergence. Loss of VDAC3 function has been associated with oxidative stress tolerance (Kanwar et al., 2022) which may explain why *vdac3-1* and *vdac3-2* did not exhibit sensitivity to submergence stress and indeed perform better during recovery. Interestingly, a recent study identified VDAC1, VDAC2 and VDAC3 as mitophagy receptors, with corresponding mutants displaying mitophagy-associated phenotypes and being unable to clear damaged mitochondria. Overall, our results suggest that the preferential activation of UPR related pathways downstream of bZIP17/28 and IRE1 are essential contributors to the superior recovery of young leaves.

### Spatiotemporal regulation of flood tolerance

Our work provides a comprehensive description of leaf-age dependent molecular responses during submergence, identifying mechanisms underlying age-specific leaf resilience and underscores the importance of the recovery phase. Plant resilience to environmental stresses is a complex interplay between developmental state and external changes. Additional complexity is conferred by the dynamic developmental progression and communication between different organs. For a leaf, such developmental changes might be linked to distinct physical and biochemical properties like leaf shape or antioxidant concentrations impacting stress tolerance. These are harder to change from a breeding perspective as this might impact plant performance also under non-stressed conditions. From the perspective of improving stress tolerance, while general resilience mechanisms have been identified (e.g. SUB1A rice (Xu et al., 2006)), in some cases more targeted strategies might be desirable. Leaf-age specific mechanisms as described here provide important insights into the spatiotemporal finetuning of stress responses and also molecular pathways and regulatory components that might form important targets for spatiotemporal manipulation of crop traits.

## Methods

### Plant material and stress treatments

Arabidopsis seeds of *bzip60/28* (Ruberti et al., 2018), *bzip28* (SALK_132285) and *bzip60* (SALK_050203), were donated by Dr. Federica Brandizzi from Michigan State University. Seeds of *vdac1* (SALK_034653), *vdac2* (SAIL_342_012), *vdac3-1* (SAIL_238_A01), *vdac3-2* (SALK_127899) and *vdac2 vdac 3-2* were donated by Prof Byung-Ho Kang and Dr Wenlong Ma from the Chinese University of Hong Kong (Ma et al., 2025). Seeds were stratified in 0.1 % agar (about 1 mg of seeds per 1 ml of 0.1 % agar) at 4°C in the dark for four days. After stratification, 40 µL of seeds in 0.1 % agar were sowed onto soil. Pots with seeds were germinated and grown in a growth chamber (20°C, 200 μmol m^-2^ s^-1^ photosynthetically active radiation (PAR), a 9-hour photoperiod from 8:00 to 17:00, and 70% relative humidity). Pots were thinned when seedlings were at 2-leaf stage (with 2 true leaves and 2 cotyledons), so there was only one seedling per pot. Plants were grown to the 10-leaf stage before they were used for experiments. Flooding treatments were as described previously (Yeung et al., 2018; Rankenberg et al., 2024). Plants were completely submerged in the dark at 20 C for the indicated durations and left to recover for the indicated time in the original climate chamber.

### RNA-seq

**For the Col-0 mRNAseq**, leaf blades of leaf 3 (old leaf) and leaf 7 (young leaf) were harvested before submergence, after 2 and 4 days of submergence, and after 1, 3, 6, and 24 hours of recovery (Fig. 1). 8-16 leaf blades were pooled together per biological replicate. For the Col-0 and *bzip28/60* RNA-seq, Arabidopsis plants were harvested before submergence, after 1 and 4 days of submergence, and after 3 hours of recovery in the light. Old (leaves 1-5) and young (leaves 6-10 and the meristem) halves of the rosettes were harvested separately. Two plants were pooled together per biological replicate. 3 biological replicates were used per genotype, tissue type and time point.

Biological replicates were harvested independently. For both RNAseq experiments, RNA was isolated using the RNeasy Plant Mini Kit (Qiagen), residual genomic DNA was digested using on-column DNase-I digestion (Ambion). RNA was quantified using a Nanodrop spectrophotometer, quality was analyzed by measuring A260/A280 and A230/A280 values on a Nanodrop and by running 1µl of RNA on an agarose gel. Library preparation was performed commercially by Macrogen Korea using the TruSeq Stranded mRNA LT Sample Prep Kit. The cDNA libraries were sequenced by paired-end Illumina sequencing on an Illumina Novaseq6000 platform, yielding approximately 30 million paired-end 150bp reads per library. Illumina sequencing of cDNA libraries from mRNA isolated from harvested samples resulted in between 30 million and 47 million high-quality paired end reads per library (Supplementary Fig. S17 and S18). These reads were then trimmed of adapter sequences and aligned to the Araport11 transcriptome yielding high mapping percentages (Supplementary Fig. S17 and S18). The mapping percentage was notably lower for Col-0 young leaves harvested after six days of submergence. A brief analysis of the origin of reads that did not map to Arabidopsis using BLAST revealed that they originated from multiple oomycetes, some viruses, as well as some unidentified sources. This suggests that the harvested Arabidopsis plants were not infected by one specific virus, as is sometimes observed in RNAseq datasets (Verhoeven et al., 2023).

### Read alignment and DEG calculation

FASTQ files were cleaned up using cutadapt (Martin, 2011) to remove some residual sequencing primers, and the quality was assessed with FastQC (Babraham Bioinformatics). Cleaned up FASTQ files were aligned to the Arabidopsis Araport11 transcriptome using Kallisto (Bray et al., 2016) on the Utrecht Bioinformatics Cluster. Differential gene expression was calculated in R using Bioconductor packages edgeR and limma (Robinson et al., 2009; Ritchie et al., 2015). Only transcripts that were present at a minimum of 20 reads in three or more samples were included and different transcript isoforms encoded from the same gene were merged. Genes were determined to be differentially expressed when they had an absolute log_2_ fold change above 1 and an FDR-corrected p-value below 0.05.

### Gene ontology and transcription factor enrichment

Gene Ontology enrichment was calculated using the R package GOseq (Young et al., 2010). Enrichment of transcription factor targets was done using the PlantRegMap database (Tian et al., 2020). Enrichment of the targets of each transcription factor per cluster was tested by hypergeometric test, adjusted for multiple testing. For visualization, transcription factors were grouped together based on their family and the percentage of overrepresented transcription factors per family was shown to account for differences in transcription factor family sizes.

### Water content

Rosettes were detached from the root system either before submergence or after the indicated submergence duration, rosettes cut off after submergence were gently blotted dry with a paper towel. Rosettes were kept on petri dishes at a relative humidity of approximately 55% and a temperature of 21°C and were weighed on a Sartorius MX-5 scale at the indicated timepoints. After the last timepoint the plants were left at 80°C for two days to dry out. Leaves were labelled as dehydrated when their water content (defined as (fresh weight – dry weight) / fresh weight * 100%) was lower than 90% of the mean water content of control leaves. A reduction of relative water content of 10% or more is typically associated with a stressful water deficit (Zhang and Bartels, 2018). Another indication of the higher water loss of old leaves was their greater reduction in leaf fresh weight (Fig. 3B). Here, leaves with a fresh weight below 40% of the mean of non-submerged leaves were designated as stressed, as this was the lower boundary of the variation in fresh weight of untreated plants.

### Stomatal aperture

Adaxial sides of leaves were pressed into President dental paste at the indicated timepoints. Imprints were made of the adaxial side, as the abaxial side of the leaf was close to the wet soil and variation in humidity across the leaf microclimate can affect stomatal behaviour (Pantin et al., 2012). After the imprints had hardened, leaves were removed and clear nail polish was applied to the imprints. Dried nail polish was mounted on microscope slides and was imaged on a Zeiss Axioskop II macroscope with a 100X magnification setting. Stomatal aperture was determined based on these pictures by measuring the ratio between the length and width of the stomatal pore using ImageJ (NIH).

### ABA treatment

ABA was applied to plants by dissolving ABA in ethanol and dissolving that in distilled water to a final concentration of 50µM or 100µM ABA, 0.1% Tween-20 was added to facilitate uptake of ABA. The mock solution contained 0.1% ethanol and 0.1% Tween-20 in distilled water. Plants recovering in the light from submergence in darkness were sprayed (200µl per spray) with either the ABA solutions or the mock twice directly after desubmergence, once 30 minutes after desubmergence, and once 1.5h after desubmergence. Stomatal aperture was determined using stomatal imprints three hours after desubmergence.

### Toluidine blue staining

Old and young leaves were cut off at the indicated timepoints, and two pooled leaves of each were submerged in either 0.05% (w/v) Toluidine Blue solution or water for two minutes. Plants were washed three times in tap water to rinse off excess staining solution and were incubated for four hours in 80% ethanol at room temperature in the dark to let all absorbed toluidine blue leach out of the leaf. 200µl of each ethanol solution was then measured for its absorbance at 626nm in a Biotek HT plate reader. Leaves were dried for 48 hours at 80°C and were weighed to normalize absorbance at 626nm by dry weight.

### Chlorophyll content

Chlorophyll extraction was performed in the dark. Each rosette was placed in a 2 ml tube and added with 1 ml DMSO. Samples were incubated at 65°C shaking bath for 30 minutes, cooled down on ice for 1 minute, and left at room temperature. Absorbance of 664 nm, 647 nm and 750 nm were measured using a spectrophotometer plate reader (Synergy HT Multi-Detection Microplate Reader). Concentrations of chlorophyll a and chlorophyll b were calculated using the equations from (Porra et al., 1989)and normalized by dry weight. Tubes containing rosettes were dried at 80°C for one week. The dry weight was measured with a microbalance scale.

### Submergence phenotype scoring

New leaf formation and dead leaf counting was done following desubmergence at different time- points during recovery under control growth conditions. New leaves were counted by marking the youngest leaf following desubmergence and then monitoring the emergence of newer leaves during recovery. Leaves were scored as dead when approximately more than half of the leaf lamina had dessicated.

### qRT-PCR

Each sample includes 2.5 μl 2X SybrGreen Supermix, 0.25 μl of 10 μM forward primer, 0.25 μl of 10 μM reverse primer and 2 μl of 5 ng/μl cDNA. The qRT-PCR was conducted on Applied Biosystems ViiA 7 Real-Time PCR System (Thermo Fisher Scientific). Forward and reverse primers of target genes we monitored are shown in Supplementary Table S6. Relative mRNA abundance was calculated by the 2^-ΔΔCT^ method as presented by Livak & Schmittgen, 2001.

### Proteomics

Arabidopsis plants Col-0 & *bzip28/60* were sampled before submergence, after 1 and 4 d of submergence, and after 3 h of recovery in the light. Young rosettes (leaves 6-10 and the meristem) were harvested. Three plants were pooled together per replicate and 4 replicates were harvested. Protein extraction, quantification, reduction, and alkalization were done as in Zhang et al. (2015). Protein precipitation was done by the methanol/chloroform method as in Zhang, et al. (2018) and sequential trypsin digestion performed at a protease: protein ratio of from 1:100 to final 1:50 (w/w). Peptide concentration was determined using a Pierce Quantitative Colorimetric Peptide Assay kit (23,275, Thermo Scientific). Labelling was performed by mixing 100 µL of peptide (1 µg/µL) with 40 µL (∼0.25mg) TMTpro™ stock in acetonitrile (TMTpro™ 16plex Mass Tag Isobaric Labelling Reagent Set Thermo Fisher Scientific) and then incubated for 1 h at room temperature.

This was followed by quenching with 8 µL of 5% hydroxylamine solution and incubation at room temperature for 15 min. Equal amounts of each prepared peptide were combined in a new tube. The solvent composition was adjusted to acetonitrile <5 % and 1 % trifluoracetic acid (HPLC grade 99.5% 044630.AE Alfa Aesar) for desalting with Sep-Pak® vac tC18 cartridge [100 mg] (Waters).

#### High-pH first dimension reverse-phase fractionation

The following LC conditions were used for the fractionation of the TMT samples: desalted peptides were resuspended in 0.1 mL 20 mM ammonium formate (pH 10.0) + 4 % (v/v) acetonitrile. Peptides were loaded onto an Acquity bridged ethyl hybrid C18 UPLC column (Waters; 2.1 mm i.d. x 150 mm, 1.7 µm particle size), and profiled with a linear gradient of 5-60% acetonitrile + 20 mM ammonium formate (pH 10.0) over 60 min, at a flow-rate of 0.25 mL/min. Chromatographic performance was monitored by sampling eluate with a diode array detector (Acquity UPLC, Waters) scanning between wavelengths of 200 and 400 nm. Samples were collected in 1 min increments and reduced to dryness by vacuum centrifugation.

#### LC-MS/MS

Dried fractions from the high pH reverse-phase separations were resuspended in 0.1% formic acid and placed into a glass vial. 1 μL of each fraction was injected by the HPLC autosampler and separated by the LC method detailed below. Fractions were combined into pairs and were analysed by LC-MS/MS.

LC-MS/MS experiments were performed using a Dionex Ultimate 3000 RSLC nanoUPLC (Thermo Fisher Scientific Inc, Waltham, MA, USA) system and a Orbitrap Lumos™ Tribrid™ mass spectrometer (Thermo Fisher Scientific Inc, Waltham, MA, USA). Peptides were loaded onto a pre-column (Thermo Scientific PepMap 100 C18, 5 mm particle size, 100 Å pore size, 300 mm i.d. x 5 mm length) from the Ultimate 3000 auto-sampler with 0.1 % formic acid for 3 min at a flow rate of 10 μL/min. After this period, the column valve was switched to allow elution of peptides from the pre-column onto the analytical column. Separation of peptides was performed by C18 reverse-phase chromatography at a flow rate of 300 nL/min using a Thermo Scientific reverse- phase nano Easy-spray column (Thermo Scientific PepMap C18, 2mm particle size, 100 Å pore size, 75 mm i.d. x 50 cm length). Solvent A was water + 0.1 % formic acid and solvent B was 80 % acetonitrile, 20 % water + 0.1 % formic acid. The linear gradient employed was 2-40 % B in 93 min. (Total LC run time was 120 min, including a high organic wash step and column re- equilibration).

The eluted peptides from the C18 column LC eluant were sprayed into the mass spectrometer by means of an Easy-Spray source (Thermo Fisher Scientific Inc.). All m/z values of eluting peptide ions were measured in an Orbitrap mass analyzer, set at a resolution of 120,000 and were scanned between m/z 380-1500 Da. Data dependent MS/MS scans (Top Speed) were employed to automatically isolate and fragment precursor ions by collision-induced dissociation [CID, Normalised Collision Energy (NCE): 35 %] which were analysed in the linear ion trap. Singly charged ions and ions with unassigned charge states were excluded from being selected for MS/MS and a dynamic exclusion window of 70 s was employed. The top 10 most abundant fragment ions from each MS/MS event were then selected for a further stage of fragmentation by Synchronous Precursor Selection (SPS) MS3 (1) in the HCD high energy collision cell using HCD [High energy Collisional Dissociation, (NCE: 55 %)]. The m/z values and relative abundances of each reporter ion and all fragments (mass range from 100-500 Da) in each MS3 step were measured in the Orbitrap analyser, which was set at a resolution of 50,000. This was performed in cycles of 10 MS3 events before the Lumos instrument reverted to scanning the m/z ratios of the intact peptide ions and the cycle continued.

Raw data were searched against the Arabidopsis TAIR10; cRAP (version of the data) database using MASCOT version 2.7.0 (Matrix Science, London, UK) and PROTEOME DISCOVERERTM v. 2.5.0.400 as described previously employing Top 10 peaks filter node and percolator nodes and reporter ions quantifier with Trypsin enzyme specificity with a maximum of one missed cleavage. Carbamidomethylation (+57.021 Da) of cysteine and TMTpro™ isobaric labelling (+304.207) of lysine and N-termini were set as static modifications while methionine oxidation (+15.996) was considered dynamic. Mass tolerances were set to 10 ppm for MS and 0.8 Da for MS/MS. For quantification, TMTpro method was used, and integration tolerance was set to Xmmu, Integration Method was set as centroid sum. Purity correction factor was set according the TMTpro™ product sheet. Each reporting ion was divided by the sum of total ions and normalized by medians of each sample. Log transformation ensured a homogeneity and normal distribution of the variances. Only proteins represented by two or more peptides were considered for further analysis. Heatmaps were generated by applying a z-score transformation using normalized log2 transformed abundance before performing the hierarchical clustering. Differential expression of proteins was determined using limma (Ritchie et al., 2015). Proteins were designated as differentially expressed when their FDR-adjusted p-value was below 0.05 and their absolute log2 fold change was above 1.

## Data availability

mRNAseq data have been deposited at the European Nucleotide Archive under accession number PRJEB57289 (Col-0) and ArrayExpress accession E-MTAB-15812 (Col-0 vs *bzip* mutants). MS proteomics data have been deposited to the ProteomeXchange Consortium via the PRIDE partner repository (Perez-Riverol et al., 2022) with the dataset identifier PXD067068.

## Acknowledgements

We thank Chelsea Rundle for assistance with proteomics data visualisation and Byung-Ho and Wenlong Ma (Chinese University of Hong Kong) for providing the *vdac* mutant seeds. Work at Rothamsted Research was funded by the Biotechnology and Biological Sciences Research Council (BBSRC) through the 21 ROMITIGATION FUND (BB/W510543/1) and the Green Engineering Institute Strategic Programme Grant (BB/X010988/1). Mass spectrometry was done at the Cambridge Centre for Proteomics. Work in the Sasidharan lab is funded by grants from the Dutch Research Council NWO (ALWOP.419; OCENW.M20.197; 016.Vidi.171.006; OCENW.M.24.182)

## Author Contributions

TR, MAS, HZ, HACFL, MBD, JRC, HVV performed the experiments and analyzed the data and revised the manuscript. TR, MAS, HZ, FLT, RS designed the experiments. TR, MAS, FLT and RS wrote the manuscript. All authors have read and approved the manuscript.

## References

1. Arias MC, Pelletier S, Hilliou F, Wattebled F, Renou JP, D’Hulst C (2014) From dusk till dawn: The arabidopsis thaliana sugar starving responsive network. Front Plant Sci. doi: 10.3389/fpls.2014.00482

2. Bray NL, Pimentel H, Melsted P, Pachter L (2016) Near-optimal probabilistic RNA-seq quantification. Nat Biotechnol. doi: 10.1038/nbt.3519

3. Bui LT, Shukla V, Giorgi FM, Trivellini A, Perata P, Licausi F, Giuntoli B (2020) Differential submergence tolerance between juvenile and adult Arabidopsis plants involves the ANAC017 transcription factor. Plant Journal. doi: 10.1111/tpj.14975

4. Cao MJ, Zhang YL, Liu X, Huang H, Zhou XE, Wang WL, Zeng A, Zhao CZ, Si T, Du J, et al (2017) Combining chemical and genetic approaches to increase drought resistance in plants. Nat Commun. doi: 10.1038/s41467-017-01239-3

5. Chang R, Jang CJH, Branco-Price C, Nghiem P, Bailey-Serres J (2012) Transient MPK6 activation in response to oxygen deprivation and reoxygenation is mediated by mitochondria and aids seedling survival in Arabidopsis. Plant Mol Biol. doi: 10.1007/s11103-011-9850-5

6. Cho HY, Chou MY, Ho HY, Chen WC, Shih MC (2022) Ethylene modulates translation dynamics in Arabidopsis under submergence via GCN2 and EIN2. Sci Adv. doi: 10.1126/sciadv.abm7863

7. D’alessandro S, Ksas B, Havaux M (2018) Decoding β-cyclocitral-mediated retrograde signaling reveals the role of a detoxification response in plant tolerance to photooxidative stress. Plant Cell. doi: 10.1105/tpc.18.00578

8. Fuchs P, Bohle F, Lichtenauer S, Ugalde JM, Feitosa Araujo E, Mansuroglu B, Ruberti C, Wagner S, Muller-Schussele SJ, Meyer AJ, et al (2022) Reductive stress triggers ANAC017-mediated retrograde signaling to safeguard the endoplasmic reticulum by boosting mitochondrial respiratory capacity. Plant Cell 34: 1375

9. Fuchs P, Rugen N, Carrie C, Elsässer M, Finkemeier I, Giese J, Hildebrandt TM, Kühn K, Maurino VG, Ruberti C, et al (2020) Single organelle function and organization as estimated from Arabidopsis mitochondrial proteomics. Plant Journal. doi: 10.1111/tpj.14534

10. Gayral M, Elmorjani K, Dalgalarrondo M, Balzergue SM, Pateyron S, Morel MH, Brunet S, Linossier L, Delluc C, Bakan B, et al (2017) Responses to hypoxia and endoplasmic reticulum stress discriminate the development of vitreous and floury endosperms of conventional maize (Zea mays) inbred lines. Front Plant Sci 8: 253326

11. Geldhof B, Novák O, Van De Poel B (2024) Leaf ontogeny modulates epinasty through shifts in hormone dynamics during waterlogging in tomato. J Exp Bot. doi: 10.1093/jxb/erad432

12. Gibbs DJ, Theodoulou FL, Bailey-Serres J (2024) Primed to persevere: Hypoxia regulation from epigenome to protein accumulation in plants. Plant Physiol 197: 584

13. Giuntoli B, Lee SC, Licausi F, Kosmacz M, Oosumi T, van Dongen JT, Bailey-Serres J, Perata P (2014) A Trihelix DNA Binding Protein Counterbalances Hypoxia-Responsive Transcriptional Activation in Arabidopsis. PLoS Biol 12: e1001950

14. Giuntoli B, Shukla V, Maggiorelli F, Giorgi FM, Lombardi L, Perata P, Licausi F (2017) Age-dependent regulation of ERF-VII transcription factor activity in Arabidopsis thaliana. Plant Cell Environ. doi: 10.1111/pce.13037

15. Hemono M, Ubrig É, Azeredo K, Salinas-Giegé T, Drouard L, Duchêne AM (2020) Arabidopsis Voltage-Dependent Anion Channels (VDACs): Overlapping and Specific Functions in Mitochondria. Cells. doi: 10.3390/cells9041023

16. Jibran R, Hunter DA, Dijkwel PP (2013) Hormonal regulation of leaf senescence through integration of developmental and stress signals. Plant Mol Biol. doi: 10.1007/s11103-013-0043-2

17. Jin HK, Hye RW, Kim J, Pyung OL, In CL, Seung HC, Hwang D, Hong GN (2009) Trifurcate feed-forward regulation of age-dependent cell death involving miR164 in Arabidopsis. Science (1979). doi: 10.1126/science.1166386

18. Jung HS, Crisp PA, Estavillo GM, Cole B, Hong F, Mockler TC, Pogson BJ, Chory J (2013) Subset of heat- shock transcription factors required for the early response of Arabidopsis to excess light. Proc Natl Acad Sci U S A. doi: 10.1073/pnas.1311632110

19. Juntawong P, Girke T, Bazin J, Bailey-Serres J (2014) Translational dynamics revealed by genome-wide profiling of ribosome footprints in Arabidopsis. Proc Natl Acad Sci U S A 111: E203–12

20. Kamal NM, Gorafi YSA, Abdelrahman M, Abdellatef E, Tsujimoto H (2019) Stay-green trait: A prospective approach for yield potential, and drought and heat stress adaptation in globally important cereals. Int J Mol Sci. doi: 10.3390/ijms20235837

21. Kanojia A, Gupta S, Benina M, Fernie AR, Mueller-Roeber B, Gechev T, Dijkwel PP (2020) Developmentally controlled changes during Arabidopsis leaf development indicate causes for loss of stress tolerance with age. J Exp Bot. doi: 10.1093/JXB/ERAA347

22. Kanwar P, Sanyal SK, Mahiwal S, Ravi B, Kaur K, Fernandes JL, Yadav AK, Tokas I, Srivastava AK, Suprasanna P, et al (2022) CIPK9 targets VDAC3 and modulates oxidative stress responses in Arabidopsis. Plant Journal. doi: 10.1111/tpj.15572

23. Khan K, Tran HC, Mansuroglu B, Önsell P, Buratti S, Schwarzländer M, Costa A, Rasmusson AG, Van Aken O (2024) Mitochondria-derived reactive oxygen species are the likely primary trigger of mitochondrial retrograde signaling in Arabidopsis. Current Biology. doi: 10.1016/j.cub.2023.12.005

24. Ko DK, Brandizzi F (2022) Transcriptional competition shapes proteotoxic ER stress resolution. Nat Plants. doi: 10.1038/s41477-022-01150-w

25. Li Z, Tang J, Srivastava R, Bassham DC, Howell SH (2020) The transcription factor bZIP60 links the unfolded protein response to the heat stress response in maize. Plant Cell. doi: 10.1105/TPC.20.00260

26. Liu M, Zwiazek JJ (2022) Transcriptomic Analysis of Distal Parts of Roots Reveals Potentially Important Mechanisms Contributing to Limited Flooding Tolerance of Canola (Brassica napus) Plants. Int J Mol Sci. doi: 10.3390/ijms232415469

27. Livak KJ, Schmittgen TD (2001) Analysis of Relative Gene Expression Data Using Real-Time Quantitative PCR and the 2−ΔΔCT Method. Methods 25: 402–408

28. Ma W, Ma J, Zhang K, Zheng X, Wang P, Feng L, Ming S, Zhuang X, Zhou J, Gao C, et al (2025) Voltage- dependent anion channels are mitophagy receptors mediating the recycling of depolarized mitochondria in Arabidopsis. bioRxiv 2025.05.29.656919

29. Müller JT, van Veen H, Bartylla MM, Akman M, Pedersen O, Sun P, Schuurink RC, Takeuchi J, Todoroki Y, Weig AR, et al (2021) Keeping the shoot above water – submergence triggers antithetical growth responses in stems and petioles of watercress (Nasturtium officinale). New Phytologist. doi: 10.1111/nph.16350

30. Mustroph A, Boamfa EI, Laarhoven LJJ, Harren FJM, Albrecht G, Grimm B (2006) Organ-specific analysis of the anaerobic primary metabolism in rice and wheat seedlings. I: Dark ethanol production is dominated by the shoots. Planta 225: 103–114

31. Mustroph A, Zanetti ME, Jang CJH, Holtan HE, Repetti PP, Galbraith DW, Girke T, Bailey-Serres J (2009) Profiling translatomes of discrete cell populations resolves altered cellular priorities during hypoxia in Arabidopsis. Proc Natl Acad Sci U S A. doi: 10.1073/pnas.0906131106

32. Ozgur R, Uzilday B, Iwata Y, Koizumi N, Turkan I (2018) Interplay between the unfolded protein response and reactive oxygen species: A dynamic duo. J Exp Bot. doi: 10.1093/jxb/ery040

33. Pantin F, Simonneau T, Muller B (2012) Coming of leaf age: Control of growth by hydraulics and metabolics during leaf ontogeny. New Phytologist. doi: 10.1111/j.1469-8137.2012.04273.x

34. Perez-Riverol Y, Bai J, Bandla C, García-Seisdedos D, Hewapathirana S, Kamatchinathan S, Kundu DJ, Prakash A, Frericks-Zipper A, Eisenacher M, et al (2022) The PRIDE database resources in 2022: a hub for mass spectrometry-based proteomics evidences. Nucleic Acids Res 50: D543–D552

35. Porra RJ, Thompson WA, Kriedemann PE (1989) Determination of accurate extinction coefficients and simultaneous equations for assaying chlorophylls a and b extracted with four different solvents: verification of the concentration of chlorophyll standards by atomic absorption spectroscopy. BBA - Bioenergetics. doi: 10.1016/S0005-2728(89)80347-0

36. Qiu K, Li Z, Yang Z, Chen J, Wu S, Zhu X, Gao S, Gao J, Ren G, Kuai B, et al (2015) EIN3 and ORE1 Accelerate Degreening during Ethylene-Mediated Leaf Senescence by Directly Activating Chlorophyll Catabolic Genes in Arabidopsis. PLoS Genet 11: e1005399

37. Rajhi I, Yamauchi T, Takahashi H, Nishiuchi S, Shiono K, Watanabe R, Mliki A, Nagamura Y, Tsutsumi N, Nishizawa NK, et al (2011) Identification of genes expressed in maize root cortical cells during lysigenous aerenchyma formation using laser microdissection and microarray analyses. New Phytologist. doi: 10.1111/j.1469-8137.2010.03535.x

38. Rankenberg T, Geldhof B, van Veen H, Holsteens K, Van de Poel B, Sasidharan R (2021) Age-Dependent Abiotic Stress Resilience in Plants. Trends Plant Sci. doi: 10.1016/j.tplants.2020.12.016

39. Rankenberg T, van Veen H, Sedaghatmehr M, Liao CY, Devaiah MB, Stouten EA, Balazadeh S, Sasidharan R (2024) Differential leaf flooding resilience in Arabidopsis thaliana is controlled by ethylene signaling-activated and age-dependent phosphorylation of ORESARA1. Plant Commun. doi: 10.1016/j.xplc.2024.100848

40. Rauniyar N, Yates JR (2014) Isobaric labeling-based relative quantification in shotgun proteomics. J Proteome Res. doi: 10.1021/pr500880b

41. Ritchie ME, Phipson B, Wu D, Hu Y, Law CW, Shi W, Smyth GK (2015) Limma powers differential expression analyses for RNA-sequencing and microarray studies. Nucleic Acids Res. doi: 10.1093/nar/gkv007

42. Robert N, d’Erfurth I, Marmagne A, Erhardt M, Allot M, Boivin K, Gissot L, Monachello D, Michaud M, Duchêne AM, et al (2012) Voltage-dependent-anion-channels (VDACs) in Arabidopsis have a dual localization in the cell but show a distinct role in mitochondria. Plant Mol Biol. doi: 10.1007/s11103-012-9874-5

43. Robinson MD, McCarthy DJ, Smyth GK (2009) edgeR: A Bioconductor package for differential expression analysis of digital gene expression data. Bioinformatics. doi: 10.1093/bioinformatics/btp616

44. Ruberti C, Lai YS, Brandizzi F (2018) Recovery from temporary endoplasmic reticulum stress in plants relies on the tissue-specific and largely independent roles of bZIP28 and bZIP60, as well as an antagonizing function of BAX-Inhibitor 1 upon the pro-adaptive signaling mediated by bZIP28. The Plant Journal 93: 155–165

45. Sachs MM, Freeling M, Okimoto R (1980) The anaerobic proteins of maize. Cell. doi: 10.1016/0092-8674(80)90322-0

46. Salinas T, Duchêne AM, Delage L, Nilsson S, Glaser E, Zaepfel M, Maréchal-Drouard L (2006) The voltage-dependent anion channel, a major component of the tRNA import machinery in plant mitochondria. Proc Natl Acad Sci U S A. doi: 10.1073/pnas.0606449103

47. Sasidharan R, Bailey-Serres J, Ashikari M, Atwell BJ, Colmer TD, Fagerstedt K, Fukao T, Geigenberger P, Hebelstrup KH, Hill RD, et al (2017) Community recommendations on terminology and procedures used in flooding and low oxygen stress research. New Phytologist. doi: 10.1111/nph.14519

48. Sasidharan R, Hartman S, Liu Z, Martopawiro S, Sajeev N, Van Veen H, Yeung E, Voesenek LACJ (2018) Signal Dynamics and Interactions during Flooding Stress. Plant Physiol 176: 1106–1117

49. Sasidharan R, Voesenek LACJ (2015) Ethylene-mediated acclimations to flooding stress. Plant Physiol. doi: 10.1104/pp.15.00387

50. Schippers JHM (2015) Transcriptional networks in leaf senescence. Curr Opin Plant Biol. doi: 10.1016/j.pbi.2015.06.018

51. Tanaka T, Tanaka H, Machida C, Watanabe M, Machida Y (2004) A new method for rapid visualization of defects in leaf cuticle reveals five intrinsic patterns of surface defects in Arabidopsis. Plant Journal. doi: 10.1046/j.1365-313X.2003.01946.x

52. Tarancón C, González-Grandío E, Oliveros JC, Nicolas M, Cubas P (2017) A conserved carbon starvation response underlies bud dormancy in woody and herbaceous species. Front Plant Sci. doi: 10.3389/fpls.2017.00788

53. Tateda C, Watanabe K, Kusano T, Takahashi Y (2011) Molecular and genetic characterization of the gene family encoding the voltage-dependent anion channel in Arabidopsis. J Exp Bot. doi: 10.1093/jxb/err113

54. Tian F, Yang DC, Meng YQ, Jin J, Gao G (2020) PlantRegMap: Charting functional regulatory maps in plants. Nucleic Acids Res. doi: 10.1093/nar/gkz1020

55. Ugalde JM, Aller I, Kudrjasova L, Schmidt RR, Schlößer M, Homagk M, Fuchs P, Lichtenauer S, Schwarzländer M, Müller-Schüssele SJ, et al (2022) Endoplasmic reticulum oxidoreductin provides resilience against reductive stress and hypoxic conditions by mediating luminal redox dynamics. Plant Cell. doi: 10.1093/plcell/koac202

56. Uno Y, Furihata T, Abe H, Yoshida R, Shinozaki K, Yamaguchi-Shinozaki K (2000) Arabidopsis basic leucine zipper transcription factors involved in an abscisic acid-dependent signal transduction pathway under drought and high-salinity conditions. Proc Natl Acad Sci U S A. doi: 10.1073/pnas.190309197

57. Van Veen H, Vashisht D, Akman M, Girke T, Mustroph A, Reinen E, Hartman S, Kooiker M, Van Tienderen P, Eric Schranz M, et al (2016) Transcriptomes of eight Arabidopsis thaliana accessions reveal core conserved, genotype- and organ-specific responses to flooding stress. Plant Physiol. doi: 10.1104/pp.16.00472

58. Verhoeven A, Kloth KJ, Kupczok A, Oymans GH, Damen J, Rijnsburger K, Jiang Z, Deelen C, Sasidharan R, van Zanten M, et al (2023) Arabidopsis latent virus 1, a comovirus widely spread in Arabidopsis thaliana collections. New Phytologist. doi: 10.1111/nph.18466

59. Voesenek LACJ, Sasidharan R (2013) Ethylene - and oxygen signalling - drive plant survival during flooding. Plant Biol. doi: 10.1111/plb.12014

60. Weits D a, Giuntoli B, Kosmacz M, Parlanti S, Hubberten H-M, Riegler H, Hoefgen R, Perata P, van Dongen JT, Licausi F (2014) Plant cysteine oxidases control the oxygen-dependent branch of the N- end-rule pathway. Nat Commun 5: 3425

61. Wenjing W, Chen Q, Singh PK, Huang Y, Pei D (2020) CRISPR/Cas9 edited HSFA6a and HSFA6b of Arabidopsis thaliana offers ABA and osmotic stress insensitivity by modulation of ROS homeostasis. Plant Signal Behav. doi: 10.1080/15592324.2020.1816321

62. Xie LJ, Tan WJ, Yang YC, Tan YF, Zhou Y, Zhou DM, Xiao S, Chen QF (2020) Long-chain acyl-CoA synthetase LACS2 contributes to submergence tolerance by modulating cuticle permeability in arabidopsis. Plants. doi: 10.3390/plants9020262

63. Xu K, Xu X, Fukao T, Canlas P, Maghirang-Rodriguez R, Heuer S, Ismail AM, Bailey-Serres J, Ronald PC, Mackill DJ (2006) Sub1A is an ethylene-response-factor-like gene that confers submergence tolerance to rice. Nature 442: 705–708

64. Yeung E, Bailey-Serres J, Sasidharan R (2019) After The Deluge: Plant Revival Post-Flooding. Trends Plant Sci. doi: 10.1016/j.tplants.2019.02.007

65. Yeung E, van Veen H, Vashisht D, Paiva ALS, Hummel M, Rankenberg T, Steffens B, Steffen-Heins A, Sauter M, de Vries M, et al (2018) A stress recovery signaling network for enhanced flooding tolerance in Arabidopsis thaliana. Proc Natl Acad Sci U S A. doi: 10.1073/pnas.1803841115

66. Young MD, Wakefield MJ, Smyth GK, Oshlack A (2010) Gene ontology analysis for RNA-seq: accounting for selection bias. Genome Biol. doi: 10.1186/gb-2010-11-2-r14

67. Yu SM, Lo SF, Ho THD (2015) Source-Sink Communication: Regulated by Hormone, Nutrient, and Stress Cross-Signaling. Trends Plant Sci. doi: 10.1016/j.tplants.2015.10.009

68. Yu WW, Chen QF, Liao K, Zhou DM, Yang YC, He M, Yu LJ, Guo DY, Xiao S, Xie RH, et al (2024) The calcium-dependent protein kinase CPK16 regulates hypoxia-induced ROS production by phosphorylating the NADPH oxidase RBOHD in Arabidopsis. Plant Cell 36: 3451–3466

69. Zagorščak M, Abdelhakim L, Rodriguez-Granados NY, Široká J, Ghatak A, Bleker C, Blejec A, Zrimec J, Novák O, Pěnčík A, et al (2025) Integration of multi-omics data and deep phenotyping provides insights into responses to single and combined abiotic stress in potato. Plant Physiol 197: kiaf126

70. Zhang H, Deery MJ, Gannon L, Powers SJ, Lilley KS, Theodoulou FL (2015) Quantitative proteomics analysis of the Arg/N-end rule pathway of targeted degradation in Arabidopsis roots. Proteomics. doi: 10.1002/pmic.201400530

71. Zhang Q, Bartels D (2018) Molecular responses to dehydration and desiccation in desiccation-tolerant angiosperm plants. J Exp Bot 69: 3211–3222

72. Zhao J, Shi M, Yu J, Guo C (2022) SPL9 mediates freezing tolerance by directly regulating the expression of CBF2 in Arabidopsis thaliana. BMC Plant Biol. doi: 10.1186/s12870-022-03445-8

73. Zhou L, Gao X, Weits DA, Zeng P, Wang X, Cai J (2021) H2S regulates low oxygen signaling via integration with the unfolded protein response in Arabidopsis thaliana. Plant Soil. doi: 10.1007/s11104-021-05091-9

