## Supplementary Figures for "Leaf-age specific flood resilience in *Arabidopsis thaliana* is determined by distinct processes contributing to post-submergence recovery"

**A)**

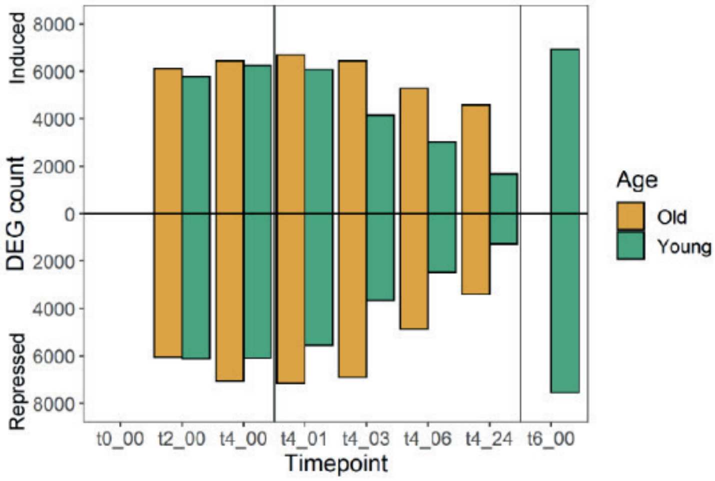


**B)**

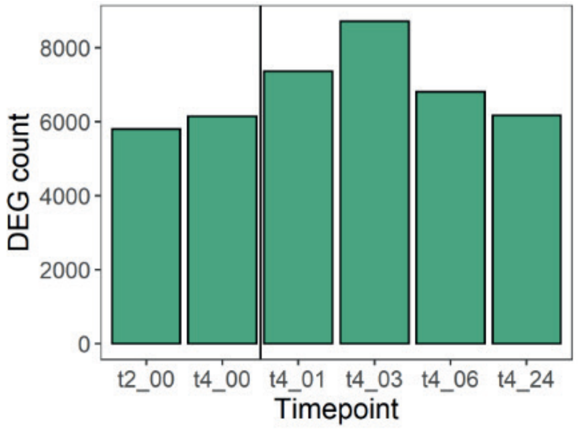


**Supplementary figure S1:**

A) DEG count for genes that responded significantly at the submergence or post- submergence timepoints compared to their non-submerged expression for old and young leaves separately. The first vertical line separates the submergence phase from the post- submergence phase. The second vertical line separates t6_00 from the samples recovering from four days of submergence.

B) DEG count for genes of which the difference in expression between old and young leaves changes significantly at each timepoint compared to non-submerged conditions. The vertical line indicates the separation between the submergence and post- submergence phases.

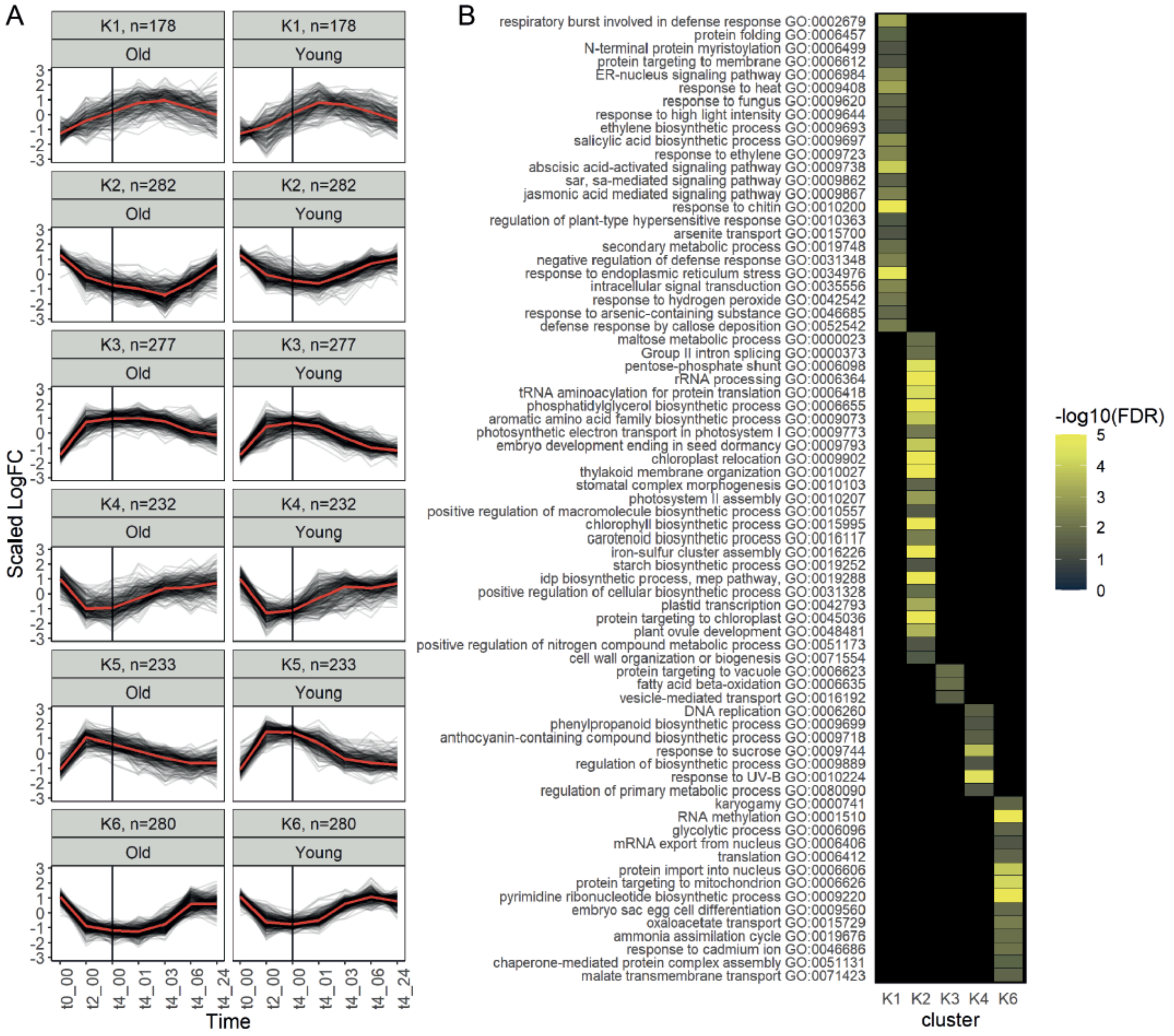


**Supplemental figure S2:** Age-independent transcriptomic responses to submergence

A) Fuzzy clustering of genes whose expression was different from non-submerged conditions in at least two samples, that never show an age dependent response. 1629 genes were divided over six clusters, yielding 1482 genes with a minimum cluster membership of 50%. Each black line represents a single gene, each red line represents the median expression in each cluster. The vertical black line indicates the separation between the submergence and the post-submergence phase.

B) Gene ontology enrichment of biological process terms in each of the six clusters shown in Figure 3A. All shown terms are significantly overrepresented in their respective cluster with an FDR-corrected p-value below 0.05.

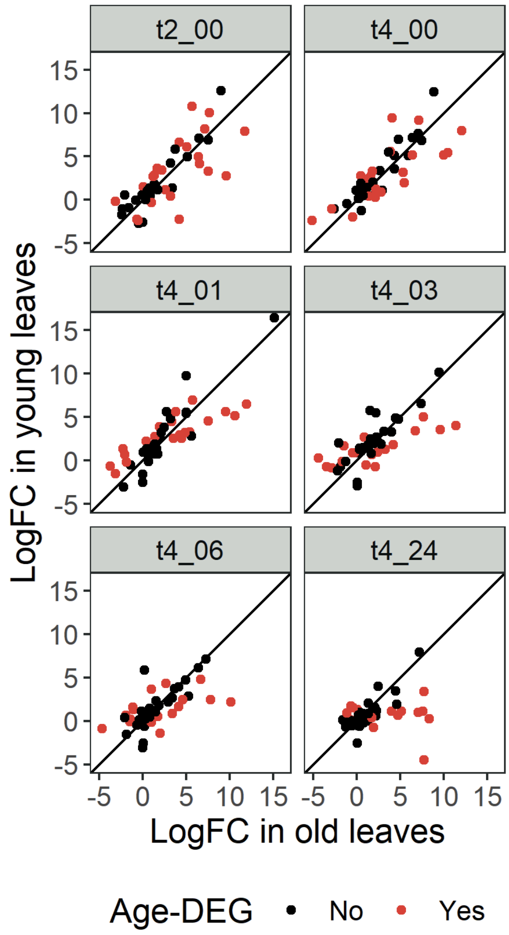


**Supplemental figure S3:** Core hypoxia genes in mRNAseq data

The response of the Arabidopsis core hypoxia gene set (Mustroph et al., 2009) to the indicated treatments. Genes with a significantly age-dependent response to the submergence or post submergence treatments are indicated in red. The diagonal line indicates where the response in old leaves (X-axis) is the same as that in young leaves (Y-axis).

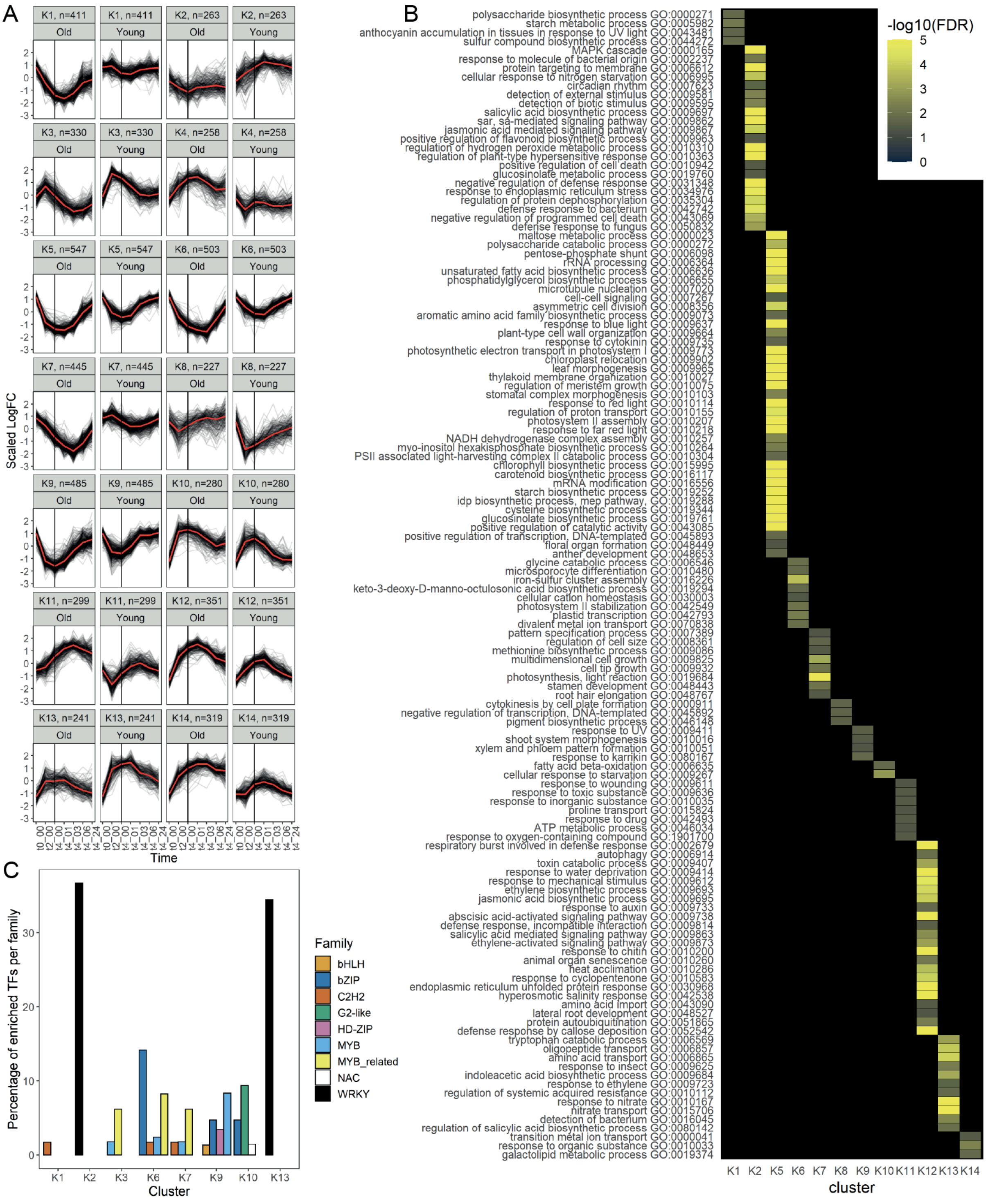


**Supplemental figure S4:** The age-dependent transcriptome response to the submergence and post submergence phase

A) Clustering of genes whose expression showed an age-dependent response at a minimum of four timepoints. 4977 genes were divided over 14 clusters, yielding 4959 genes with a minimum highest cluster membership of 20%. Each black line represents a single gene, each red line represents the median expression in each cluster. The vertical black line indicates the separation between the submergence and post-submergence phase.

B) Gene ontology enrichment of the biological process terms in each of the 14 clusters shown in Figure 4A. All shown terms were significantly overrepresented in their respective cluster with an FDR-corrected p-value below 0.05.

C) Enrichment of targets of transcription factors in each cluster, grouped by transcription factor family. Enrichment was calculated by hypergeometric tests, corrected for multiple testing. Families with fewer than 2 transcription factors with overrepresented targets were omitted.

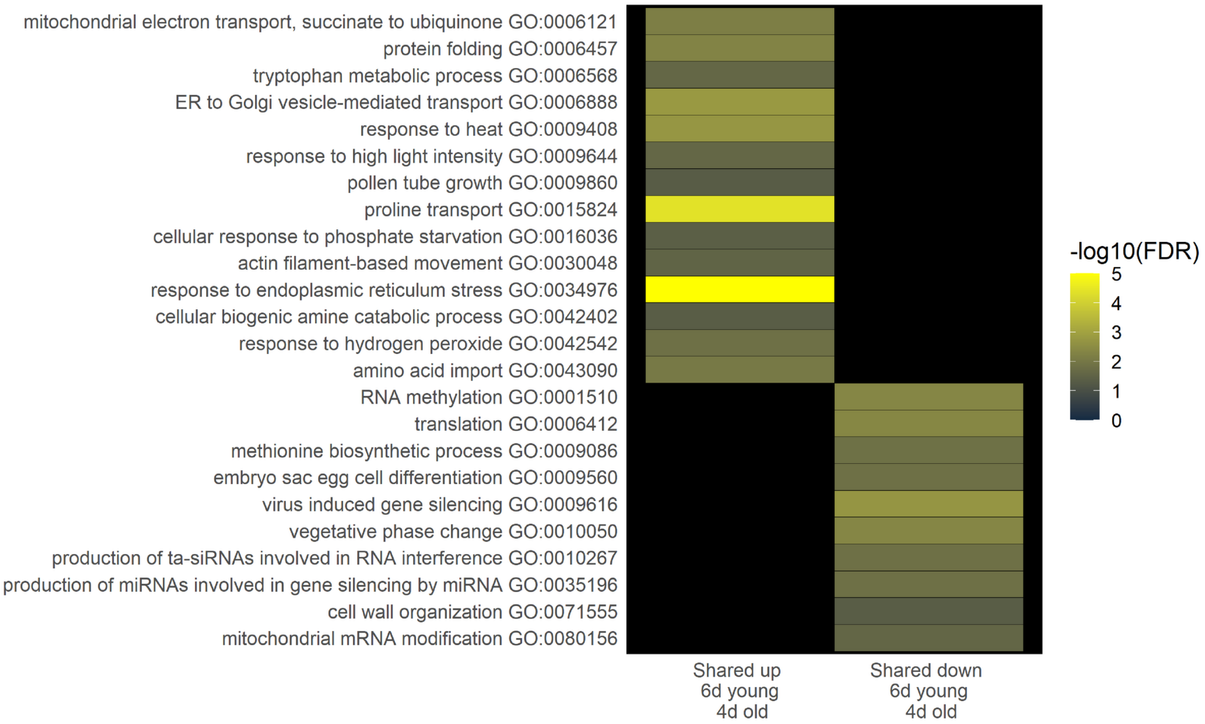


**Supplemental figure S5:** Significantly enriched biological process GO categories among genes that were up- or downregulated in old leaves after four days of submergence and in young leaves after six days of submergence. These were the last harvested submergence timepoints for old and young leaves. All shown terms were significantly overrepresented in their respective overlapping dataset with an FDR-corrected p-value below 0.05.

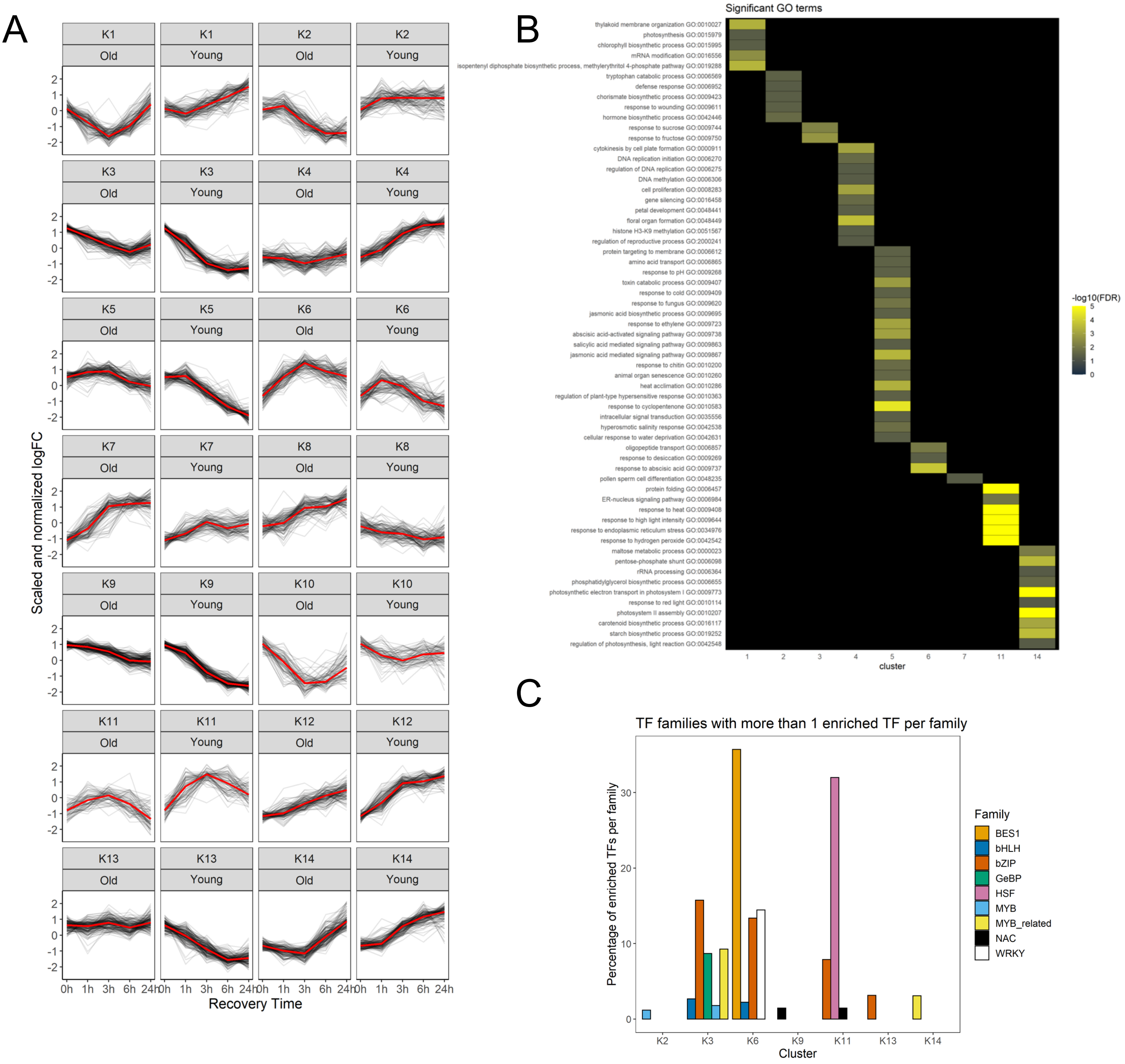


**Supplemental figure S6:** The age-dependent transcriptome response to post-submergence recovery

A) Clustering of genes whose expression shows an age-dependent response at a minimum of three recovery timepoints. 1848 genes were divided over 14 clusters, yielding 1845 genes with a minimum cluster membership of 20%. Each black line represents a single gene, each red line represents the median expression in each cluster.

B) Gene ontology enrichment of each of the 14 clusters shown in Figure 7A.

C) Enrichment of targets of transcription factors in each cluster, grouped by transcription factor family.

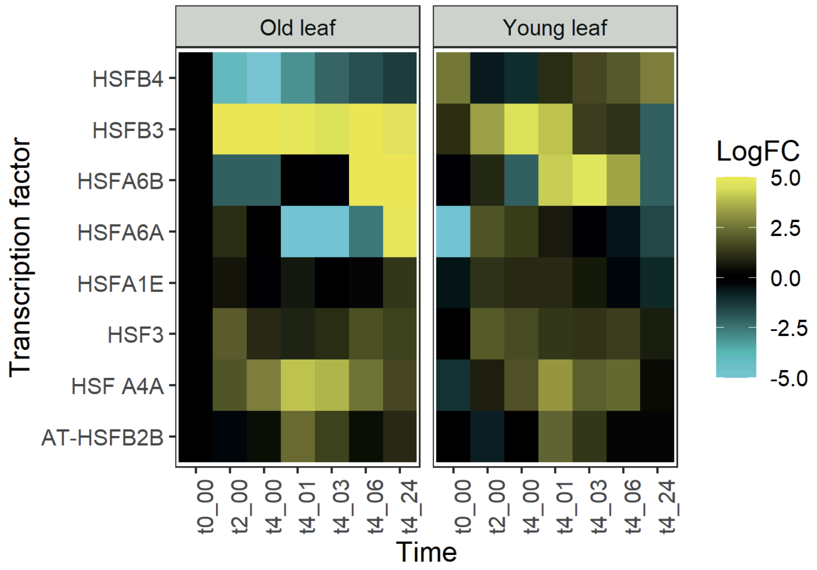


**Supplemental figure S7:** Expression of HSF transcription factors.

Expression of the eight transcription factors of the HSF family whose targets are enriched in cluster K11 (Supplementary figure S6). Expression was calculated relative to that of non-submerged old leaves.

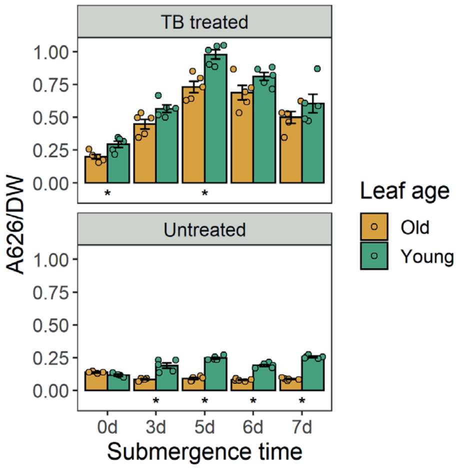


**Supplemental figure S8**: Cuticular permeability as measured by toluidine blue (TB) uptake of old and young leaves. Absorbance was measured at 626nm, values were normalized by leaf dry weight. Leaves not treated with TB were used to control for baseline differences in absorbance at 626nm. n=4-5 per group, each consisting of two old or young leaves from different plants pooled together. Asterisks indicate significant differences between old and young leaves (paired t-test).

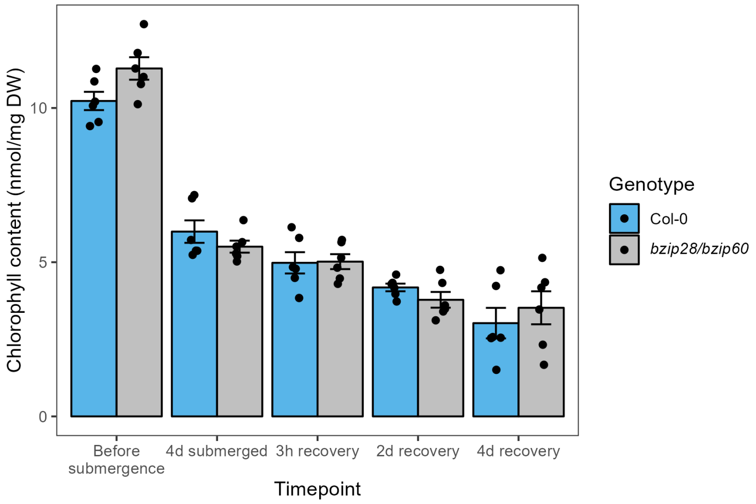


**Supplemental figure S9:** Senescence is not affected by disruption of the ER sensing pathway. Chlorophyll levels were measured for Col-0 and *bzip28/bzip60* double mutant plants submerged for 4d and then allowed to recover (3h, 2d, 4d). Data represents standard errors of the mean (SEM) where n=6. Two-way ANOVA with Tukey's multiple comparisons test was performed and found not to be significant for genotype comparisons $p<0.05$.

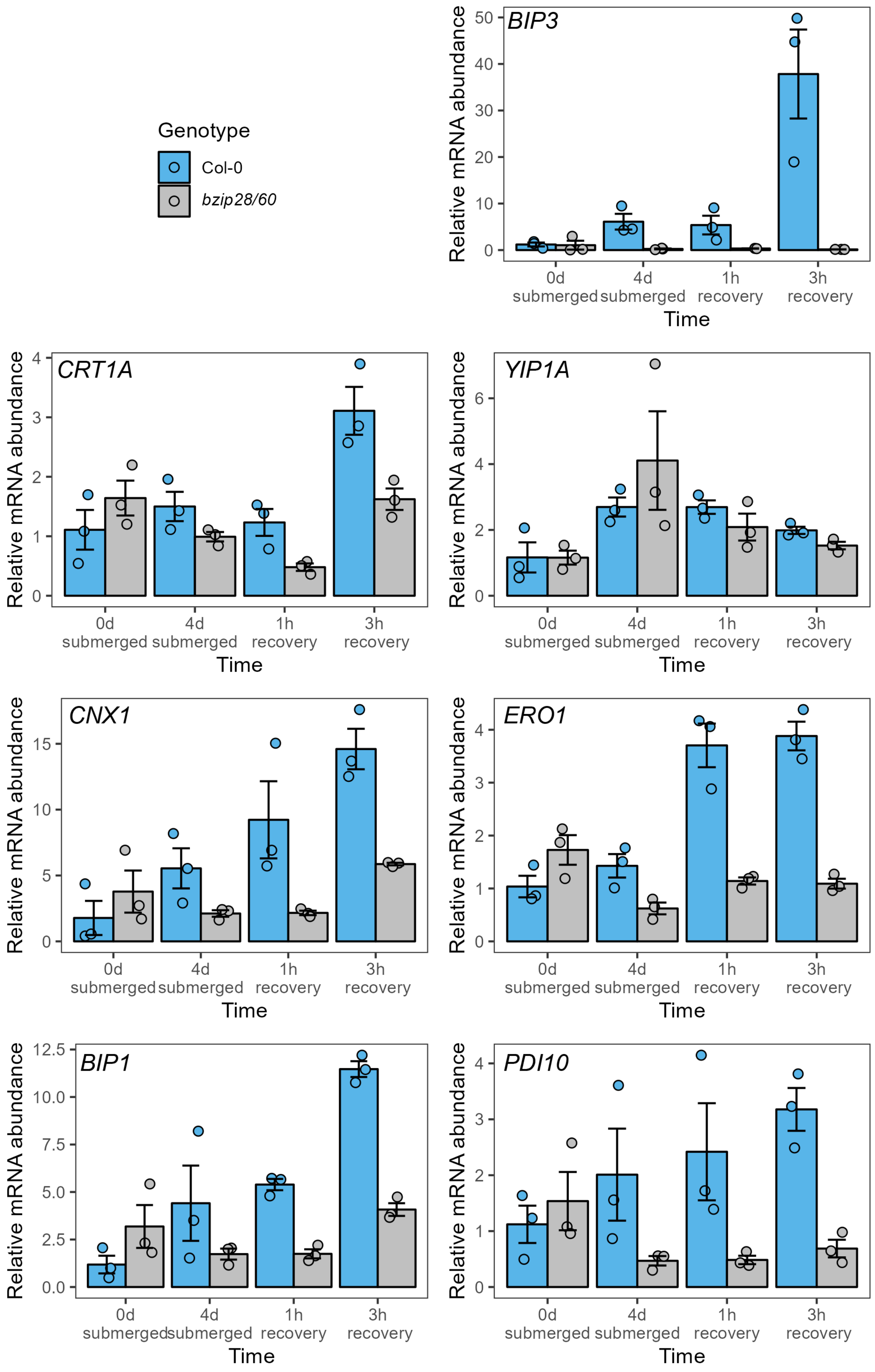
Genotype
Col-0
bzip28/60
BIP3
CRT1A
YIP1A
CNX1
ERO1
BIP1
PDI10
Relative mRNA abundance
0d submerged
4d submerged
1h recovery
3h recovery
Time

**Supplemental figure S10:** Unfolded Protein Response (UPR) genes are predominantly accumulated during the recovery phase. Plants were submerged for 4d and allowed to recover for three hours. Shoot tissue was collected from Col-0 and *bzip60/28* plants and mRNA levels were measured by qRT-PCR using the *Actin2* gene as a normalization control. Quantified genes included *BIP1, BIP3* (*Binding protein 1 and 3*), *CNX1*(*Calnexin 1*), *CRT1A* (*Calreticulin 1A*), *PDI10* (*Protein disulfide isomerase 10*) and *ERO1* (*Endoplasmic reticulum oxidoreductin1*). Data represent means with SEM in both directions among three biological replicates. Each biological replicate contains one shoot rosette.

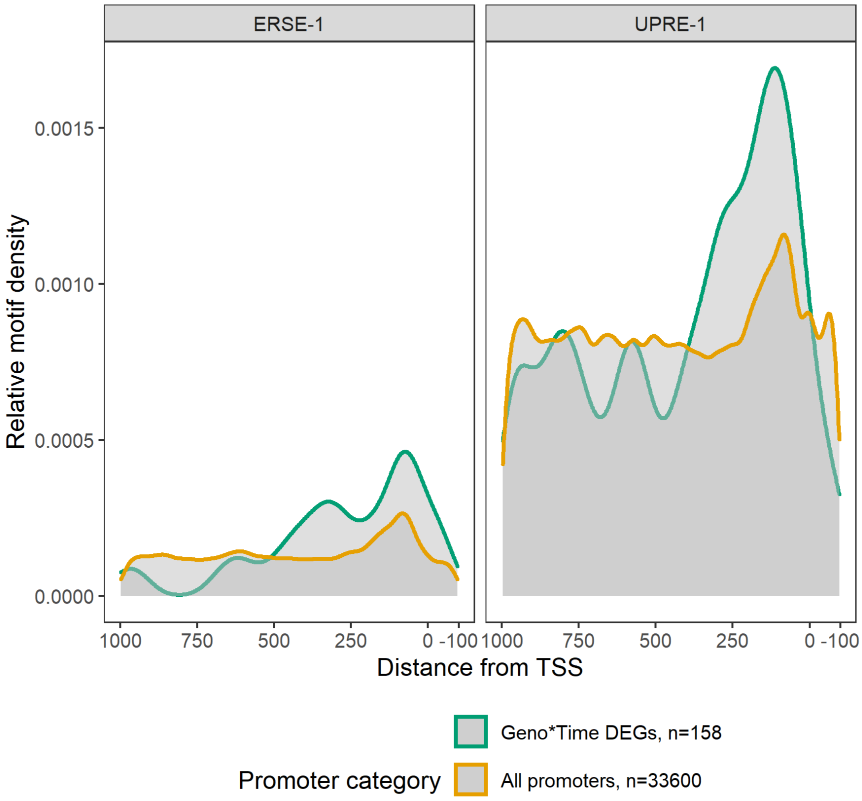


**Supplemental figure S11:** Relative density of ER-stress related transcriptional motifs ERSE-1 and UPRE-1 (Ko and Brandizzi, 2024), compared between all promoters (orange line; non-DEGs) or promoters that show a significant genotype*time interaction effect at any comparison. (green lines).

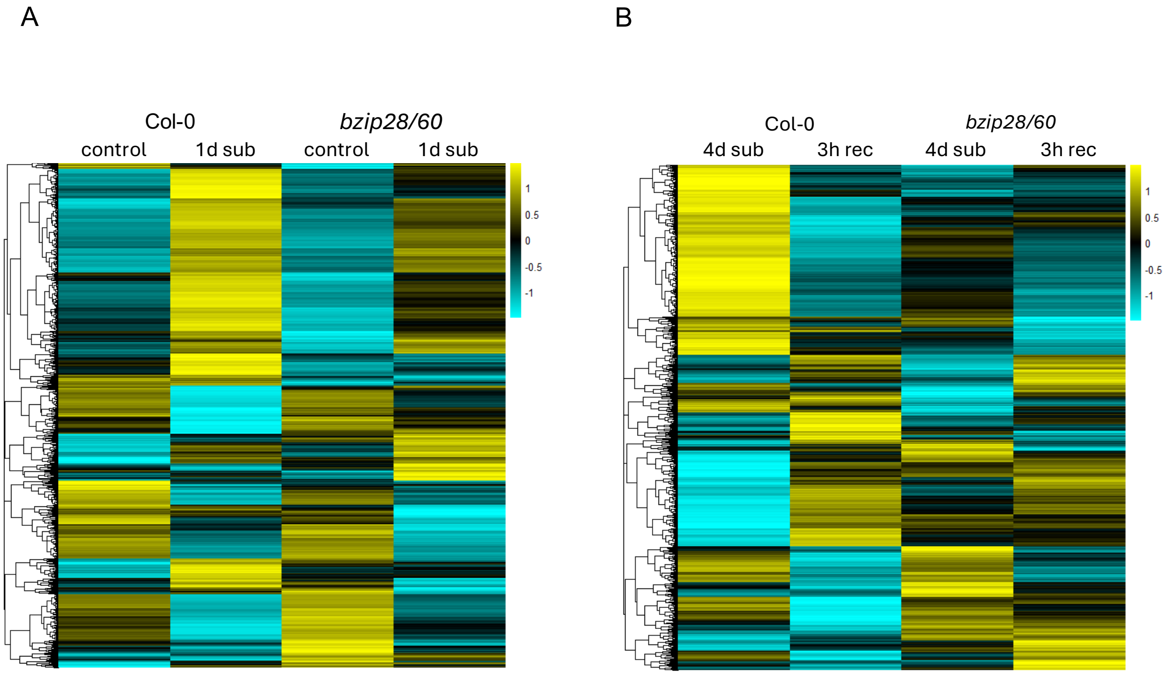


**Supplemental Figure S12:** Heatmaps of scaled protein abundance for (A) 3,608 proteins with at least 2 peptides in the 1d submergence experiment; (B) 4,767 proteins with at least 2 peptides in the 4d submergence-3 h recovery experiment.

A

B

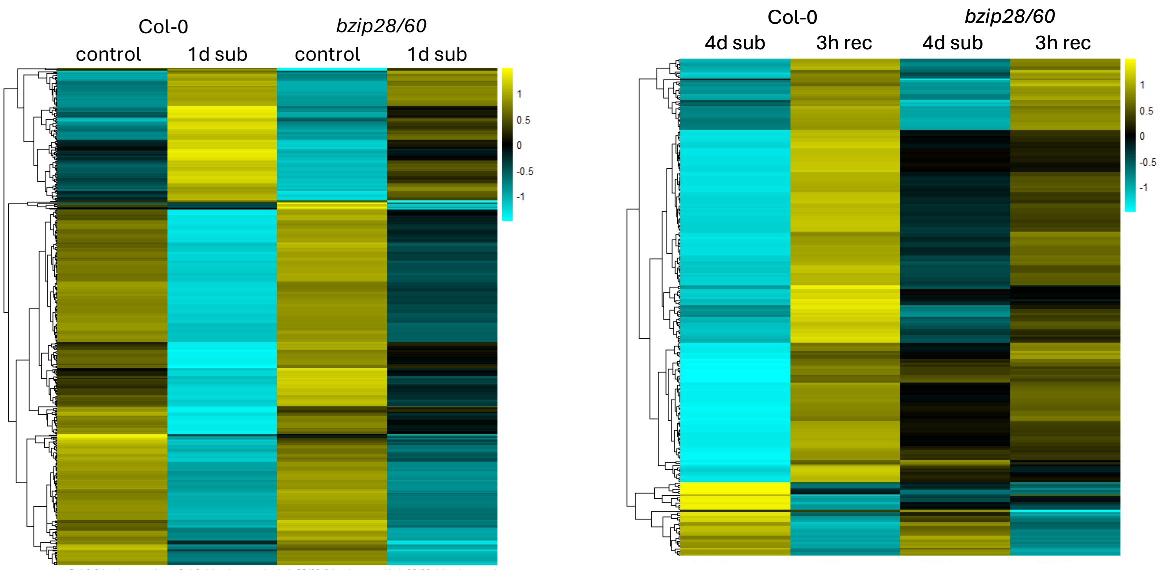


**Supplemental Figure S13:** Heatmaps of scaled protein abundance for: (A) 460 proteins with significant 1.5-fold changes ($p < 0.05$) in at least one comparison of Col-0 and *bzip28/60* in the 1d submergence experiment; (B) 250 proteins with significant 1.5-fold changes ($p < 0.05$) in at least one comparison of Col-0 and *bzip28/60* in the 4d submergence-3 h recovery experiment.

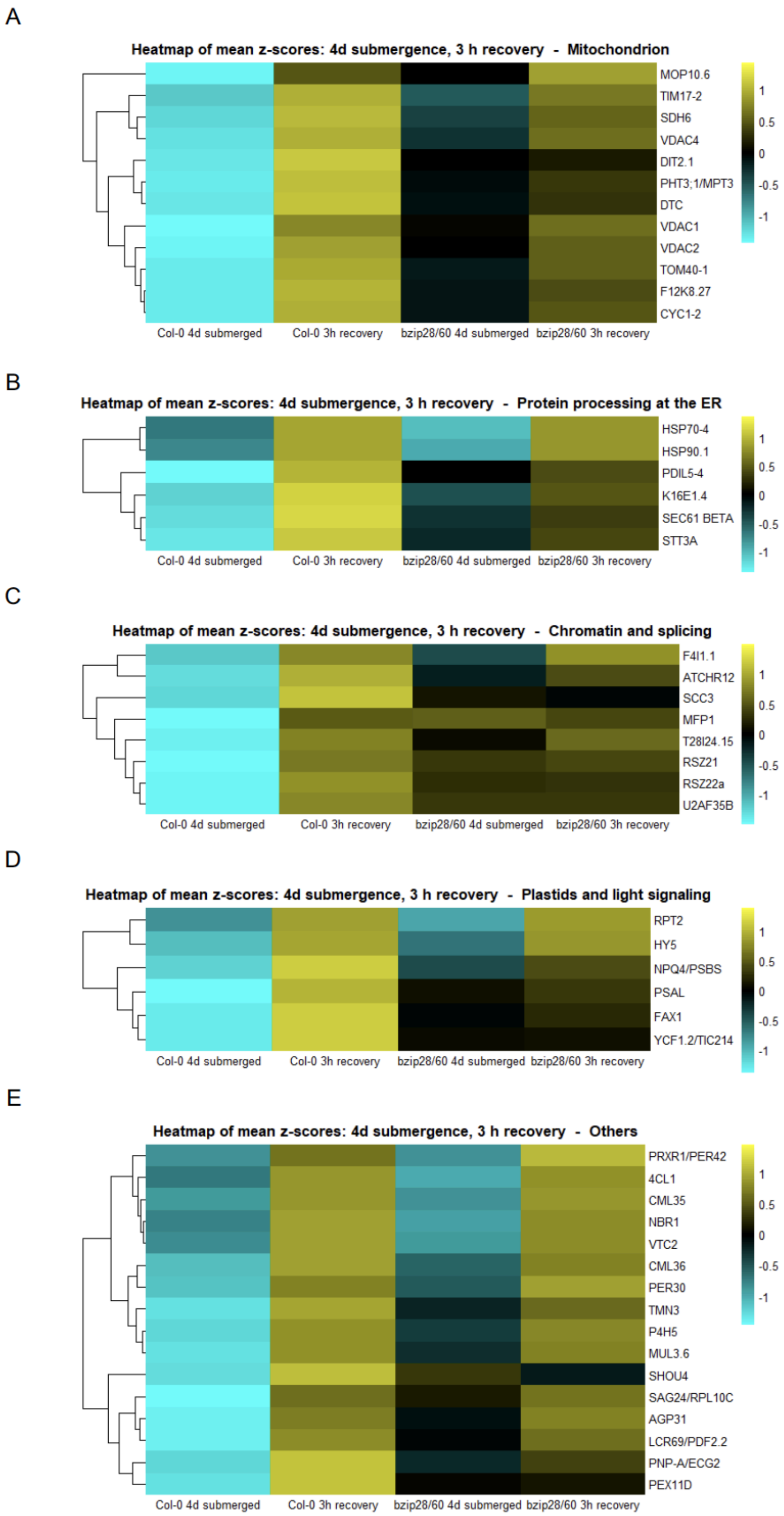
A
Heatmap of mean z-scores: 4d submergence, 3 h recovery - Mitochondrion
MOP10.6
TIM17-2
SDH6
VDAC4
DIT2.1
PHT3;1/MPT3
DTC
VDAC1
VDAC2
TOM40-1
F12K8.27
CYC1-2
Col-0 4d submerged
Col-0 3h recovery
bzip28/60 4d submerged
bzip28/60 3h recovery
1
0.5
0
-0.5
-1
B
Heatmap of mean z-scores: 4d submergence, 3 h recovery - Protein processing at the ER
HSP70-4
HSP90.1
PDIL5-4
K16E1.4
SEC61 BETA
STT3A
Col-0 4d submerged
Col-0 3h recovery
bzip28/60 4d submerged
bzip28/60 3h recovery
1
0.5
0
-0.5
-1
C
Heatmap of mean z-scores: 4d submergence, 3 h recovery - Chromatin and splicing
F4I1.1
ATCHR12
SCC3
MFP1
T28I24.15
RSZ21
RSZ22a
U2AF35B
Col-0 4d submerged
Col-0 3h recovery
bzip28/60 4d submerged
bzip28/60 3h recovery
1
0.5
0
-0.5
-1
D
Heatmap of mean z-scores: 4d submergence, 3 h recovery - Plastids and light signaling
RPT2
HY5
NPQ4/PSBS
PSAL
FAX1
YCF1.2/TIC214
Col-0 4d submerged
Col-0 3h recovery
bzip28/60 4d submerged
bzip28/60 3h recovery
1
0.5
0
-0.5
-1
E
Heatmap of mean z-scores: 4d submergence, 3 h recovery - Others
PRXR1/PER42
4CL1
CML35
NBR1
VTC2
CML36
PER30
TMN3
P4H5
MUL3.6
SHOU4
SAG24/RPL10C
AGP31
LCR69/PDF2.2
PNP-A/ECG2
PEX11D
Col-0 4d submerged
Col-0 3h recovery
bzip28/60 4d submerged
bzip28/60 3h recovery
1
0.5
0
-0.5
-1

**Supplemental Figure S14:** Heatmaps of scaled protein abundance for different categories of proteins with increased abundance in Col-0 after 3 h recovery. (A) Mitochondrial proteins; (B) Protein processing at the ER; (C) Chromatin and splicing; (D) Plastids and light signalling; (E) Others.

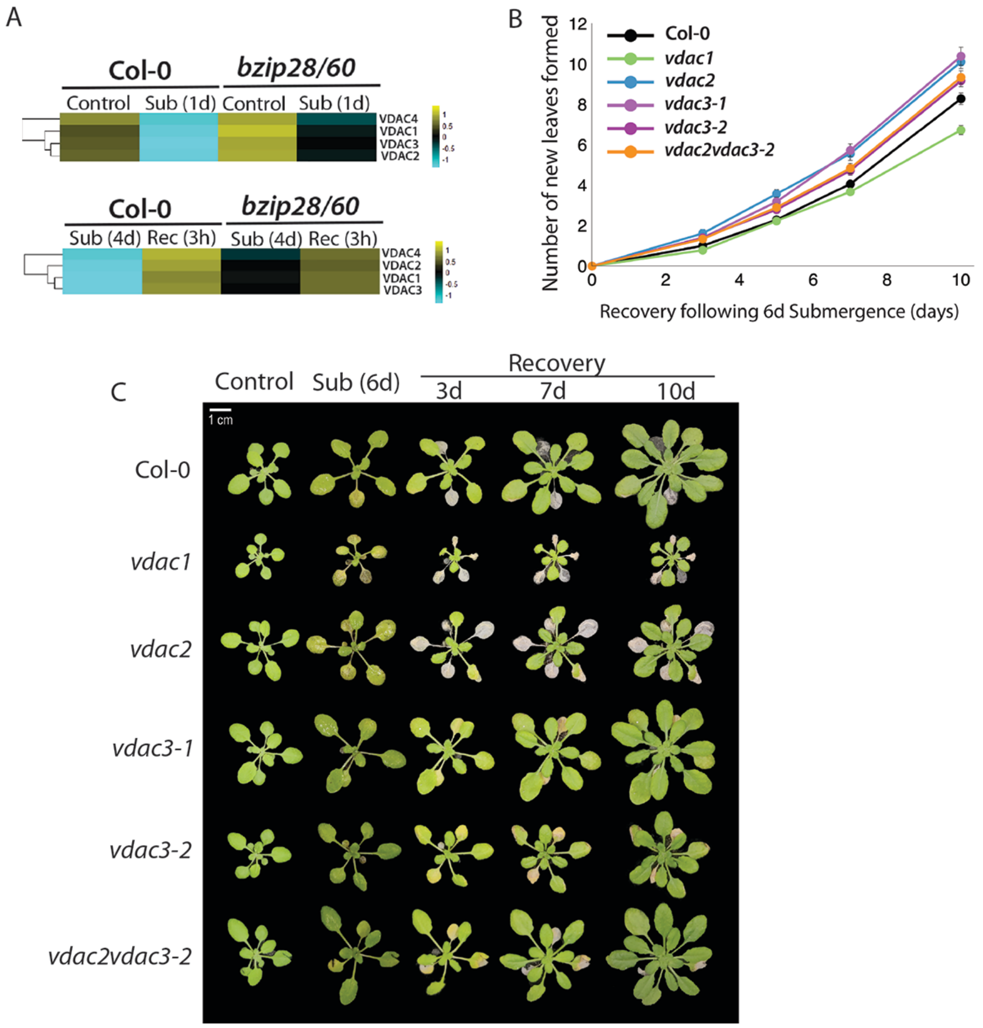


**Supplemental Figure S15:** Heatmaps of scaled protein abundance for VDAC proteins in Col-0 and *bzip28/60* mutants (A) before and after 1 d submergence (upper panel); and after 4 d submergence and 3 h post-submergence recovery (lower panel). (B) Mean number of new leaves formed in Col-0 and *vdac1, vdac2, vdac3-1, vdac3-2* mutants after 6 d submergence, followed by 3, 6, 7 d or 10 d of recovery. Statistical significance was calculated using a two-way ANOVA followed by a Tukey's post-hoc test. At the 10d recovery time-point significant differences in new leaf formation were observed for *vdac1*, *vdac2* and *vdac3-1* (See Supplemental Table S3). (C) Representative images of plants of Col-0 and *vdac* mutants after 6 d of submergence, followed by 3, 7, or 10 d recovery.

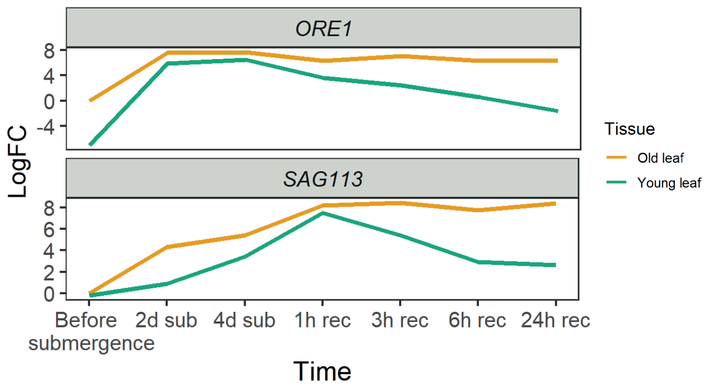


**Supplemental figure S16:** Expression of *ORE1* and *SAG113* before, during and after submergence. Expression was calculated relative to that in old leaves before submergence.

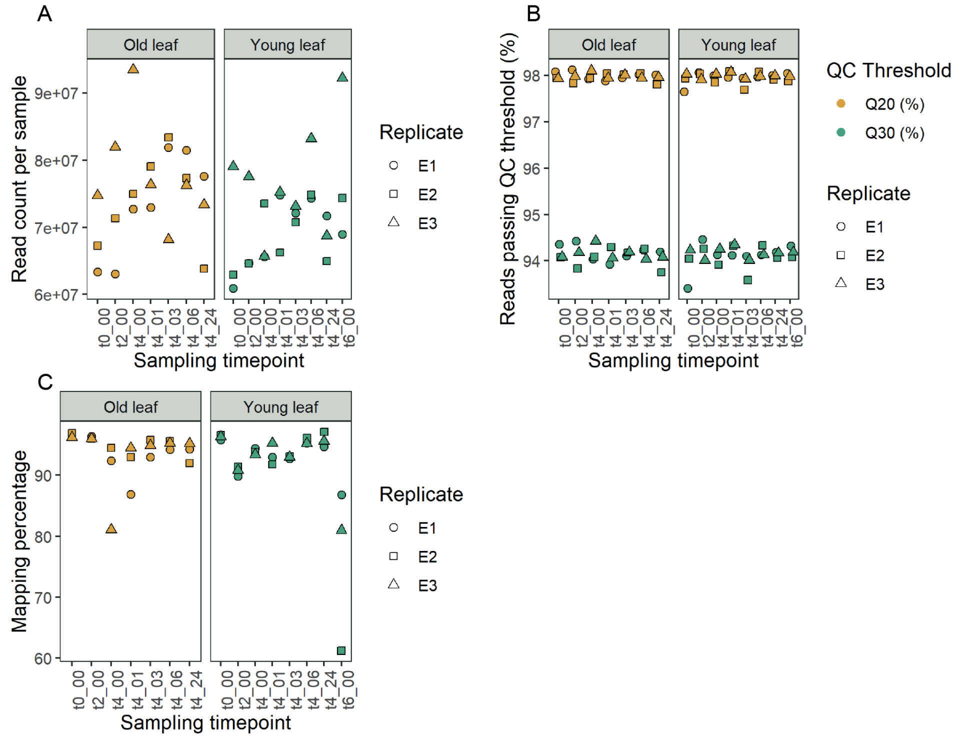


**Supplementary figure S17:** Quality control of mRNAseq (Col-0 old versus young leaves, see methods)

A) Number of sequenced reads per library of old and young leaves. Shapes indicate different biological replicates
B) Percentage of reads of each library passing the Q20 (99% accuracy) and Q30 (99.9% accuracy) Phred quality thresholds
C) Percentage of reads per library mapped to the Araport11 transcriptome.

A)

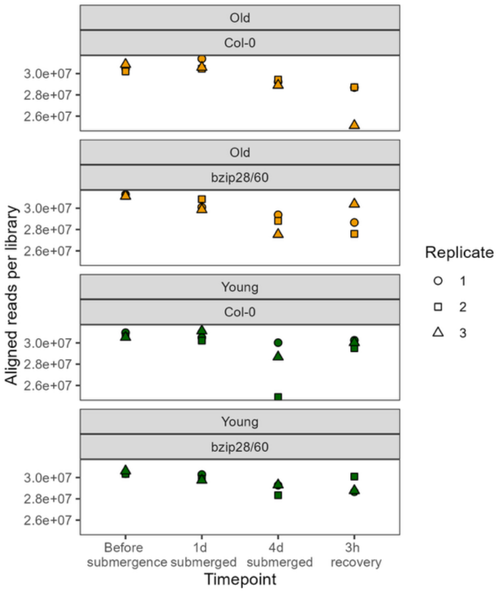


B)

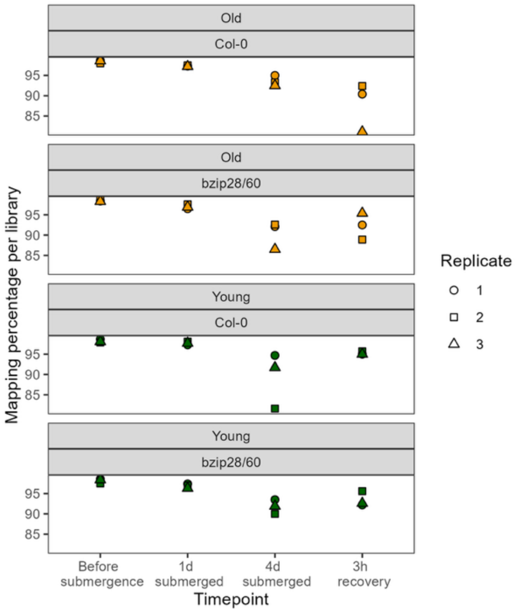


**Supplementary figure S18:** Quality control of mRNAseq (Col-0 vs *bzip28/60*, old versus young leaves, see methods)

A) Number of sequenced aligned reads per library of old and young leaves. Shapes indicate different biological replicates

B) Percentage of reads per library mapped to the Araport11 transcriptome.
